# Three-dimensional and molecular brain atlas of the hagfish reveals the evolutionary origin and early diversification of the vertebrate brain

**DOI:** 10.64898/2025.12.03.692081

**Authors:** Riho Harada, Motoki Tamura, Hirofumi Kariyayama, Shigenori Nonaka, Daichi G. Suzuki

**Author notes:** These authors contributed equally to this work (Riho Harada, Motoki Tamura).

## Abstract

Hagfish and lampreys form the clade cyclostomes, the earliest branching lineage of the vertebrates. Although hagfish are generally assumed to have experienced intense modifications, its brain development shows certain ancestral patterns that are secondarily lost in the lamprey. Therefore, hagfish provides key insights into the evolutionary origin and early diversification of the vertebrate brain. However, neuroanatomical information of the hagfish brain is substantially limited today due to its sources based on old-fashioned methodology alone. Here, we provide a detailed brain atlas of the adult hagfish, based on three-dimensional reconstruction and expression analysis of genetic markers for major neuronal transmitters. Through our analyses, we found the following key characteristics. First, the dopaminergic system appears to be distributed more broadly than previously thought in the hagfish brain, implying that dopamine is involved in various neural functions. Second, the previously suggested “primordial cerebellum” area in hagfish shows notable affinity to the cerebellum-like octavolateral structures of the jawed vertebrate hindbrain. Last, the gene expression profile suggests a hippocampus-equivalent brain region in hagfish, that is, the ventrolateral subnucleus of the central prosencephalic complex (NCvl). This study highlights conserved and diversified neuroarchitecture of the hagfish brain, providing a pivotal reference for further studies.

## Introduction

Hagfish was once a “key player” in comparative neuroanatomy. Before the rise of molecular phylogenetics, this animal was situated as the sister group of the “true vertebrates” (i.e., lampreys and jawed vertebrates), forming together a larger group called craniates (the craniate theory). This theory was based on seeming morphological plesiomorphies of hagfish, such as the lack of vertebrae and de-epithelialized neural crest cells (Ota & Kuratani, 2006). In this context, the hagfish brain was vigorously investigated to elucidate the evolutionary origin of the vertebrate brain as reviewed in Wicht & Nieuwenhuys (1998). However, recent molecular phylogenetic studies (Blair & Hedges, 2005; Delarbre et al., 2002; Delsuc et al., 2006; Furlong & Holland, 2002; Mallatt & Sullivan, 1998; Stock & Whitt, 1992; Suga et al., 1999; Takezaki et al., 2003) consistently support the monophyly of the hagfish and the lamprey (the cyclostome theory), further confirmed by a series of studies in evolutionary developmental biology showing that the once-thought ancestral states found in hagfish are in fact secondary modifications (Higuchi et al., 2019; Ota et al., 2007, 2011). In consequence, the lamprey brain has been simply focused on as the representative model of the cyclostome or ancestral vertebrate brain. The hagfish brain indeed appears to have experienced extensive secondary specializations. For example, its lateral ventricles are obliterated and its subpallial forebrain region is occupied by enigmatic neuronal clusters called the central prosencephalic complex (Wicht & Nieuwenhuys, 1998). The peculiarities of the hagfish are often linked to the adaptation to its deep-water habitat and scavenging benthic life, as suggested by studies in comparative morphology and paleontology (Fernholm & Holmberg, 1975; Gabbott et al., 2016).

However, this attitude was challenged by Sugahara et al. (2016), showing that gene expression patterns for vertebrate brain development are strongly conserved in the hagfish. While there are, of course, hagfish-specific derivative characters such as the lack of the epiphysis, the authors demonstrate that the hagfish retain some ancestral traits missing in the lamprey. Particularly, in hagfish and jawed vertebrates, hedgehog (*Hh*) is expressed in the ventral-most part of the primordial telencephalon and marks this region as the medial ganglionic eminence (MGE), the precursor of the pallidum. Although the pallidum-equivalent region is suggested to be present in the adult lamprey brain (Stephenson-Jones et al., 2011), in contrast, no *Hh* gene expression is found in the developing telencephalon of this animal (Sugahara et al., 2011, 2016). This finding implies that lineage-specific modifications have been accumulated not only in the hagfish but also in the lampreys. Following this assumption, a paleontological study suggests that the lamprey have evolutionarily experienced a substantial change in their life cycle, into which the larval and metamorphic periods were secondarily inserted (Miyashita et al., 2021).

Essentially, at least three clades are required to infer their ancestral states. In the above-mentioned case, for example, we cannot draw conclusions about the origin of the MGE based solely on the fact that lampreys lack this structure whereas jawed vertebrates possess it. Meanwhile, recent research on hagfish brain development allows us to determine this evolutionary polarity; the lamprey shows the derivative state, while the hagfish and jawed vertebrates retain the ancestral state inherited from the common ancestor of all vertebrates (Sugahara et al., 2016). Similarly, a genomic survey of hagfish olfactory repertoire revealed that hagfish possess type 2 vomeronasal receptors (V2Rs), which had been regarded to be absent in cyclostomes as the lamprey lack them (Kariyayama et al., 2025). This finding suggests that the evolutionary origin of these receptors can be traced further back, from the common ancestor of jawed vertebrates, as previously thought, to the common ancestor of all vertebrates (Kariyayama et al., 2025). Given that both the lamprey and hagfish show ancestral and derivative characters in a mosaic manner, we thus need information from both lineages for ancestral state reconstruction.

Therefore, even in the present situation where the cyclostome theory is widely accepted, investigation of the hagfish brain is crucial in deciphering the early evolution of the vertebrate brain. Nevertheless, our understanding of the complex neural structures is currently based on old-fashioned methodologies such as histological sections and manual schematic drawings (e.g., Wicht & Northcutt 1993), leaving room for subjective interpretation. Furthermore, the homology of brain regions between the hagfish and other vertebrates remains unclear because gene expression analysis has not been performed for the adult hagfish brain yet. In this study, we thus attempted to employ modern techniques (i.e., three-dimensional digital reconstruction and gene expression profiling) to establish a state-of-the-art brain atlas of the hagfish.

## Materials and Methods

### Animals

*E. burgeri* specimens (both male and female) were collected from Sagami Bay, Kanagawa Prefecture, from October to June each year, 2021–2024. The animals were transported to the laboratory at the University of Tsukuba, Japan. They were kept in saltwater aquaria at 12 C° before dissection. All animal experiments were performed under the guidelines for animal use of the Animal Care Committee at the University of Tsukuba.

### Brain fixation and sampling

To obtain brain samples, perfusion fixation was performed. First, animals were deeply anesthetized with MS-222 (100 mg/L; Sigma, A5040), and their branchiocardial region was exposed. Next, the outlet needle of the perfusion fixation tool set (Genostaff, INT-PFTS-M) was inserted into the ventricle. Subsequently, an incision to the atrium was made, allowing blood to be expelled. By drip infusion, blood was replaced by artificial seawater containing 50 μg/ml heparine, and then by 4% paraformaldehyde (PFA)/PBS, 100 ml each. After perfusion fixation, the brain was harvested and postfixed with 4% PFA/PBS for 1 hour at room temperature (RT) or overnight at 4 C°. The fixed brain was then rinsed with PBS, dehydrated in a graded methanol series (25%, 50%, 75%, and 100%), and stored at −20 °C.

### Whole-mount fluorescent Nissl staining and three-dimensional digital reconstruction

Fixed brains were stained by NeuroTrace 530/615 Red Fluorescent Nissl Stain (Invitrogen, N21482) and clarified with the CUBIC Trial Kit (Fujifilm, 290-80801). First, the samples were rehydrated with PBS and then soaked in 50% CUBIC-1 Solution in PBS containing 0.1% Tween20 (PBST) overnight, followed by 100% CUBIC-1 Solution for two days at RT. After washing with PBST for one day, they were incubated in 5 μl/ml fluorescent Nissl reagent/PBST for four weeks at RT. After washing with PBST once again for one day, they were subsequently immersed in 50% CUBIC-2 Solution/PBST overnight, then in 100% CUBIC-2 Solution for two days at RT. The stained and clarified specimens were examined using a light-sheet microscope (Zeiss, Lightsheet Z.1).

The acquired data were first tiled using ZEN software (Zeiss, Zen blue 3.4) and then analyzed using Amira 3D (Thermo Fisher Scientific, version 2021.1). Each brain region was identified and segmented based on the density of neuronal somata stained by fluorescent Nissl reagent. Finally, the segmented regions were colored and reconstructed in three-dimensional images.

### Gene search and phylogenetic analysis

We performed BLASTP (2.15.0+; Camacho et al., 2009) searches against a previously published genome assembly of *E. burgeri* (Yu et al., 2024) on the Ensembl genome browser (Eburgeri_3.2; https://www.ensembl.org/Eptatretus_burgeri) using the amino acid sequence of the following query proteins: *Petromyzon marinus* (sea lamprey) Vesicular glutamate transporter (VGluT/Slc17a6b2, GenBank: XP_032815942), *P. marinus* Glutamate decarboxylase (GAD, GenBank: XP_032821372), *P. marinus* Serotonin transporter (SerT/Slc6a4, GenBank: XP_032802967), *P. marinus* (sea lamprey) Choline acetyltransferase (ChAT, GenBank: XP_032814983), *Mus musculus* (mouse) tyrosine hydroxylase (TH, GenBank: NP_033403), mouse dopamine transporter (DAT/Slc6a3, GenBank: NP_034150), *Myxine glutinosa* (Atlantic hagfish) D1 dopamine receptor (GenBank: CAA06542), and *Lampetra fluviatilis* (European river lamprey) D2 dopamine receptor (GenBank: ADO23655). The suggested sequences were then used as queries for searches in global protein databases at NCBI and checked for their close affinity to homologous genes of interest (reciprocal best hits).

For phylogenetic analysis, the suggested sequences were aligned using MAFFT (v.7; Katoh & Standley, 2013) and trimmed by trimAl (Capella-Gutiérrez et al., 2009). We used the automated1 option of trimAl for the *VGluT*, *ChAT*, *TH*, *GAD*, *DAT*/*NET* analysis, and the strict option for the *DRD* genes. Phylogenetic trees were generated by the maximum likelihood method using RAxML (version 8.2.12; Stamatakis, 2014) with the options of “-f a -x 12345 -p 12345 -# 500 -m PROTGAMMAAUTO --auto-prot=aic -T 16”.

### Microsynteny analysis for DAT/NET and D2 family dopamine receptors

We first collected the genome annotations of humans (*Homo sapience*), mice (*Mus musculus*), tropical clawed frogs (*Xenopus tropicalis*), zebrafish (*Danio rerio*), ghost sharks (*Callorhinchus milii*), and hagfish (*Eptatretus burgeri*) from ENSEMBL release 111 (https://ensembl.org/). The genome annotation of the sea lamprey (*Petromyzon marinus*) was obtained by the NCBI Reference Sequence Database (https://www.ncbi.nlm.nih.gov/datasets/genome/GCF_010993605.1/). We then extracted the 20 protein-coding sequences flanking *Drd2*, *DAT*, and *NET* genes from each species. To identify the genes of these obtained sequences, we retrieved orthology information from human proteome data in BioMart Ensembl. For lamprey sequences, we performed BLASTP searches against human proteome data to identify their orthology with E value < 1E-10. Based on these results,we visualized the gene order in respective species using the tidyverse package and the ggplot2 package in R version 4.2.2.

### Molecular cloning

Total RNA was extracted from whole brains of adult *E. burgeri* using TRIzol Reagent (Invitrogen, 15596026). These RNAs were reverse transcribed into cDNAs using PrimeScript II 1st strand cDNA Synthesis Kit (Takara, 6210A) and were used as templates in the polymerase chain reaction (PCR). Hagfish genes of interest were amplified by PCR using the primers listed in Table 1. We used reverse primers including the T3 promoter sequence (20 bp) to synthesize DIG-labeled RNA probes for *in situ* hybridization.

**Table 1.**
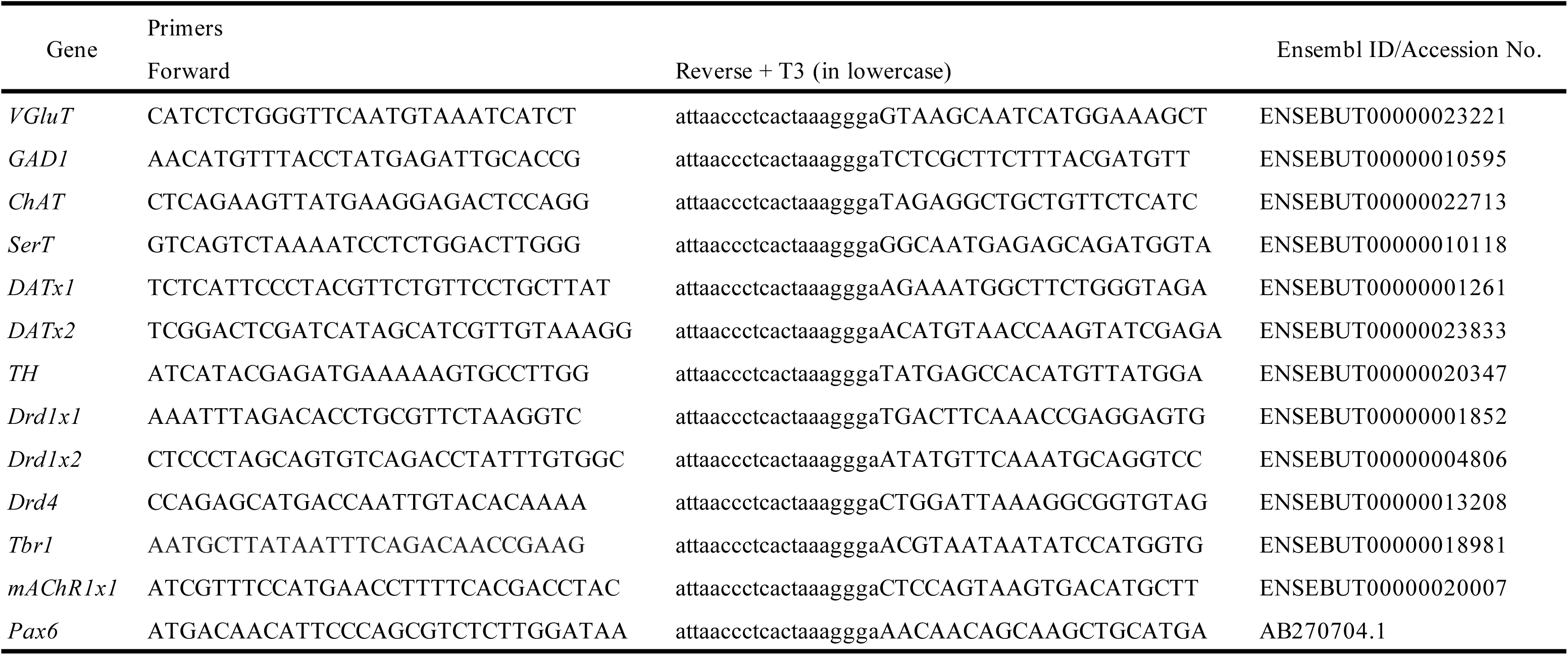
Primer sequences for RNA probes.

### Section *in situ* hybridization and fluorescent Nissl staining

Fixed brains stored in 100% methanol at −20 °C were rehydrated in a graded methanol series (75%, 50%, and 25% in PBS) and PBS. The specimens were replaced with 10% and then 30% sucrose. Subsequently, the specimens were embedded in Tissue-Tek Optimum Cutting Temperature (O.C.T.) Compound (Sakura Finetek Japan, 4583). Frozen sections in 20 μm were prepared using a cryostat (Leica, CM1860) and mounted on MAS-coated slide glasses (Matsunami Glass Ind., SMAS-01). These sections are served for *in situ* hybridization and fluorescent Nissl staining described below.

DIG-labeled antisense RNA probes were transcribed from PCR products using T3 RNA polymerase (Roche, 11031163001). After sections were washed with PBS to remove the O.C.T. compound, they were postfixed for 20 min with 4% PFA/PBS and rinsed with PBS. Later, they were prehybridized in hybridization buffer (50% Formamide, 5× SSC, 5× Denhaldt’s solution, 50 μg/ml yeast tRNA, and 50 μg/ml heparin in deionized-distilled H_2_O, hereafter ddH_2_O) and then incubated in the hybridization buffer with 0.1 mg/ml probe overnight at 60 °C. Subsequently, the specimens were washed with the following solutions: 50% Formamide and 2× SSC (for 30 min at 60 °C, twice), 2× SSC (for 30 min at 60 °C, twice), 0.2× SSC (for 30 min at 60 °C, twice), and PBS (5 min at RT). For signal detection, the specimens were blocked with 0.5% blocking reagent (Roche, 11277073910) in PBS for 1 hour at RT and then incubated with 0.5 μl/ml alkaline phosphatase (AP)-conjugated anti-digoxigenin Fab fragments (diluted in 0.5% blocking reagent/PBS; Roche, 11093274910) overnight at 4 °C. After washing with tris-buffered saline (TBS) four times for 30 min each at RT, alkaline phosphatase activity was detected with 20 μl/ml NBT/BCIP in color development buffer (100 mM Tris HCl pH 9.5, 100 mM NaCl, and 50 mM MgCl_2_). The stained specimens were postfixed in 4% PFA/PBS, dehydrated in a graded methanol series (25%, 50%, 75%, 95%, and 100%) and xylene-equivalent solvent G-NOX (Genostuff, GN04), and mounted using xylene-free mounting medium PARAmount-D (Falma, 308-500-1). The prepared slides were examined under a microscope (Nikon, ECLIPSE Ni) and photographed using a microscope digital camera (Nikon, DS-Ri1).

For fluorescent Nissl staining, after sections were washed with TBS to remove the O.C.T. compound, they were incubated in 0.1% NeuroTrace 530/615 Red Fluorescent Nissl Stain (Invitrogen, N21482)/TBS for 2 hours. They were then washed with TBS and mounted with ProLong Diamond Antifade Mountant (Invitrogen, P36961). The prepared slides were examined under a fluorescence microscope (Nikon, ECLIPSE Ni; Nikon, D-LEDI-C) and photographed using a microscope digital camera (Nikon, DS-Ri1). The acquired photographs were then monochromatized and black/white-inverted using Adobe Photoshop (version 25.7).

### Acetylcholinesterase (AChE) histochemistry

Fresh brains were harvested from *E. burgeri* adults that were deeply anesthetized with MS-222 and then euthanized by decapitation. After washing with PBS, they were put in a disposable plastic dish Tissue-Tek Cryomold (Sakura Finetek Japan, 4566) filled with the O.C.T. compound and rapidly frozen on an aluminum block at −80 °C. Frozen sections in 40 μm were prepared using a cryostat (Leica, CM1860) and mounted on MAS-coated slide glasses (Matsunami Glass Ind., SMAS-01).

AChE staining was performed according to Eilam-Altstädter et al. (2022) with minor modifications. The sections were washed with distilled water to remove the O.C.T. compound and were incubated in reaction buffer (50 mM sodium acetate buffer pH 5.0, 10 mM glycine, 0.2 mM CuSO_4_·5H_2_O, 1.156 mg/ml S-acetylthiocholine iodide, and 60 μg/ml ethopropazine in deionized-destilled H_2_O) for 24–36 hours at RT. After washing with tap water, signals were developed in 10% K-Ferricyanide in deionized-destilled H_2_O). The stained specimens were washed with PBS twice, dehydrated in a graded ethanol series (25%, 50%, 75%, 95%, and 100%) and G-NOX, and mounted using PARAmount-D. The prepared slides were examined under a microscope (Nikon, ECLIPSE Ni) and photographed using a microscope digital camera (Nikon, DS-Ri1).

## Results

### Three-dimensional neuroarchitecture of the hagfish brain

The hagfish brain is divided into five major parts (Fig. 1A); the olfactory bulb (bulbus olfactorius, BO), telencephalon (TEL), diencephalon (DIE), mesencephalon (MES), and rhombencephalon (RH). According to Wicht & Northcutt (1992) and Wicht & Nieuwenhuys (1998), we further subdivided some of these regions (Fig. 1D): the telencephalon into the pallium in five layers (P1–5), the central nucleus (nucleus centralis, NC), and the preoptic area (PO); the diencephalon into the habenula (HA), the thalamus (TH), the hypothalamus (HY), the synencephalon (SY), and the posterior tuberculum (tuberculum posterior, TP); the mesencephalon into the midbrain tectum (tectum mesencephali, TM) and the tegmentum (TEG).

**Figure 1.**
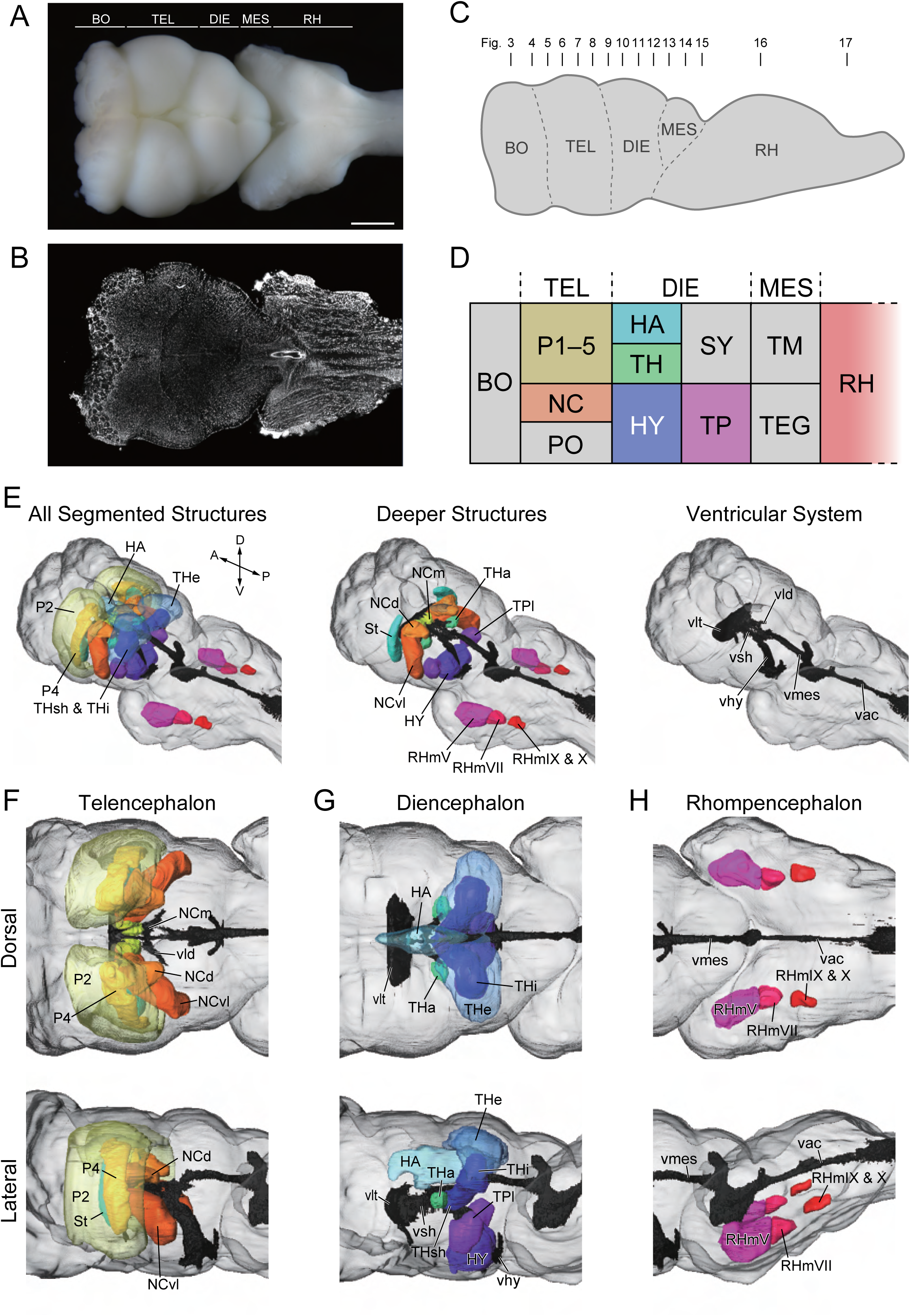
Overview and three-dimensional structures of the hagfish brain. (A) Hagfish brain in dorsal view. (B) Horizontal plane section of the whole-mount lightsheet scanning of the hagfish brain, of which neural somata are stained with fluorescent Nissl dye. (C) Schematic image of the hagfish brain in lateral view, showing the sectioning plane levels for the gene expression analysis. (D) Major brain regions of the hagfish brain in lateral view. (E) Overview of the three-dimensional model of the hagfish brain: all segmented structures (left), deeper structures (middle), and the ventricular system (right). (F–H) Three-dimensional reconstruction of the telencephalic (F), diencephalic (G), and rhombencephalic (G) brain regions. Scale bar: 1 mm in (A) for (A, B).

Tissue clarification and light-sheet microscopy enabled us to observe the inner structure of the hagfish brain (Fig. 1B). We segmented different brain regions *in silico* based on fluorescent Nissl signals of neuronal somata. Based on the obtained results, we reconstructed the shapes and relative positions of these brain regions (Fig. 1E–H).

As previously described (Wicht & Northcutt, 1992; Wicht & Nieuwenhuys, 1998), the hagfish pallium possesses five layers, of which the second (P2) and the fourth (P4) layers contain more neuronal somata. Our results indicated that the P2 distributes in a broad superficial part of the dorsal telencephalon, while the P4 occupies a relatively restricted portion (Fig. 1F). We also confirmed that there are both telencephalic and diencephalic lateral ventricles (vlt and vld) in the adult hagfish brain (Fig. 1E, F, G), although Wicht & Northcutt (1992) did not describe the latter. We also found that the dorsal and ventrolateral parts of NC (NCd and NCvl) were placed between these two ventricular parts, suggesting that these nuclei push the lateral ventricle aside and consequently divide it into two anterior telencephalic (vlt) and posterior diencephalic (vld) portions (see also Wicht & Nieuwenhuys 1998).

Based on this reconstruction, we subsequently performed molecular analysis for cross sections at different rostrocaudal levels as shown in Fig. 1C.

### Identification of neurotransmitter-marker and receptor genes

Vesicular glutamate transporter (VGluT) is a class of glutamate transporters often used as a marker for glutamatergic neurons (e.g., Villar-Cervino et al., 2011). We found one VGluT candidate in *E. burgeri*. As this homolog showed close affinity to known vertebrate VGluT sequences (Fig. 2A), we concluded it is certainly hagfish VGluT.

**Figure 2.**
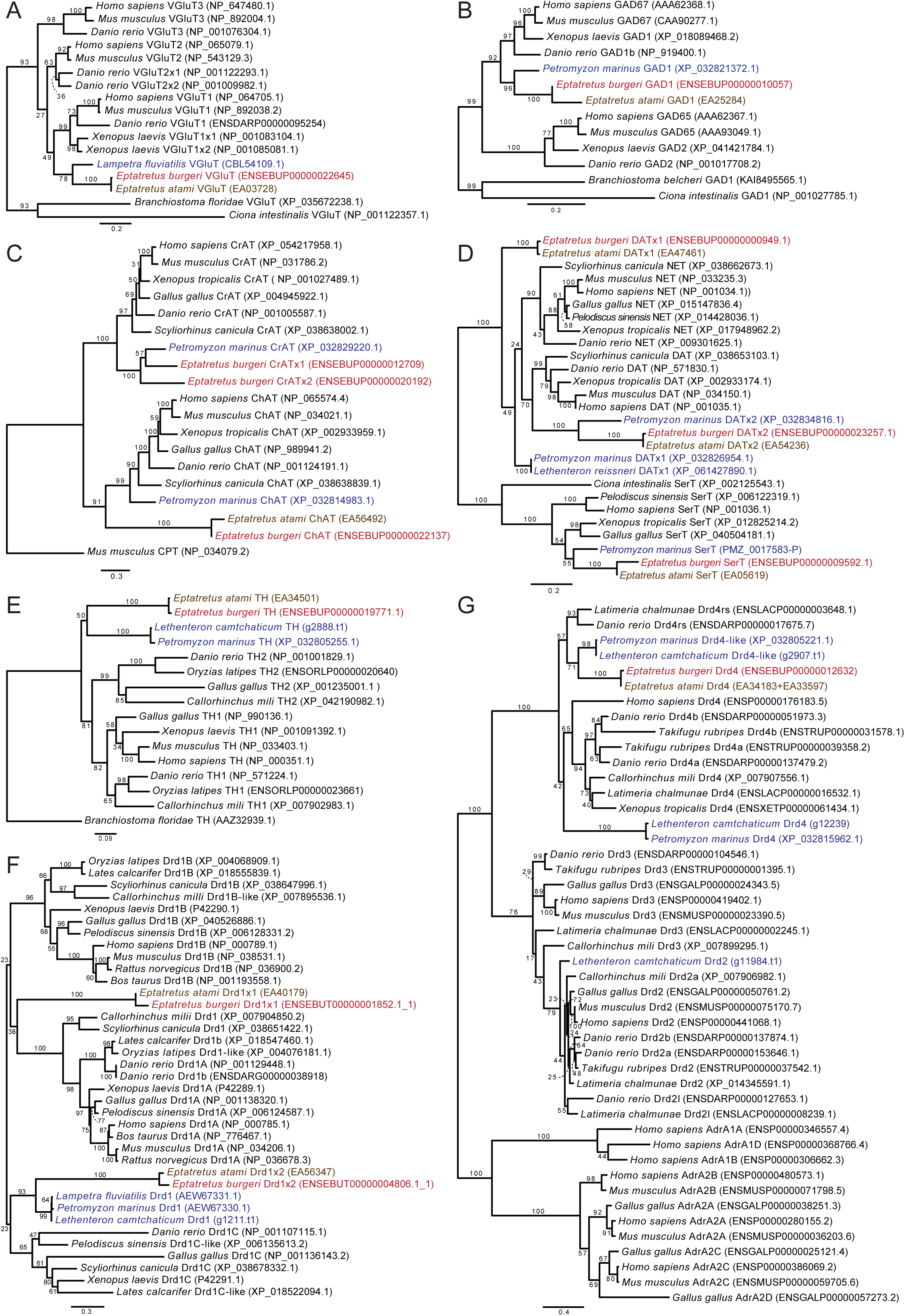
Phylogenetic trees of the major neurotransmitter markers, based on amino acid sequences. (A) VGluT, (B) GAD, (C) ChAT, (D) DAT/NET/SerT, (E) TH, (F) Drd1 (D1), (G) Drd2/3/4. The numbers described on some nodes indicate exact bootstrap values. Scale bar represents the number of amino acid substitutions per site.

Glutamate decarboxylase (GAD) is required for γ-aminobutyric acid (GABA) production in GABAergic neurons (Ueno, 2000). In vertebrates, *GAD* genes are divided into two major subfamilies, GAD1/GAD67 and GAD2/GAD65 (Bu & Tobin, 1994; Lariviere et al., 2002). In hagfish, we found one GAD sequence, which formed a sister group with its lamprey homolog in GAD1 subfamily (Fig. 2B). Based on this result, we defined this homolog as hagfish GAD1.

Choline acetyltransferase (ChAT) is responsible for acetylcholine synthesis in cholinergic neurons (Oda, 1999). We found that hagfish have one type of ChAT as the basal-most branch of all vertebrate ChAT homologs (Fig. 2C). We also found two other *E. burgeri* genes similar to known vertebrate ChAT, but they actually formed a cluster with carnitine acetyltransferase (CrAT) as an outgroup of ChAT.

Serotonin transporter (SerT) is commonly used as a serotonergic neuron marker (Blakely et al., 1994), while dopamine transporter (DAT) is expressed in dopaminergic neurons (Nepal et al., 2023). DAT, SerT, and norepinephrine transporter (NET) are collectively categorized as monoamine transporters (Aggarwal & Mortensen, 2017). We found one candidate for SerT and two for DAT in hagfish. Our phylogenetic analysis indicated that the candidate gene for SerT indeed belongs to the SerT subfamily (Fig. 2D) One of the candidate genes for hagfish DAT (hagfish DATx2), together with a lamprey DAT candidate, formed a sister group of gnathostome DAT genes, while the other (hagfish DATx1) was located in a lineage basal to all vertebrate DAT and NET sequences (Fig. 2D). To assess the gene homology, we further conducted synteny analysis (Suppl. Fig. 1). As a result, we found that hagfish DATx1 and DATx2 were in line with Lpcat1 and Lpcat2 respectively. In mice, DAT and NET are found with lysophosphatidylcholine acyltransferase 1 (Lpcat1) on chromosome 13 and lysophosphatidylcholine acyltransferase 2 (Lpcat2) on chromosome 8, respectively. These results suggest that hagfish DATx1 is orthologous to mouse DAT, while DATx2 corresponds to mouse NET. This orthology was also supported by expression analysis described below.

Tyrosine hydroxylase (TH) is known as a marker gene for dopaminergic neurons, involved in dopamine synthesis (Daubner et al., 2011). We found one hagfish TH homolog and confirmed that it forms a sister group with its lamprey homolog in the TH subfamily (Fig. 2E).

In mammals, five subtypes of dopamine receptors are known and divided into two distinct families (Yamamoto et al., 2015); the D1 family consists of two subtypes (D1 and D5), while the D2 family comprises the rest (D2, D3, and D4). We found two D1 family homologs and one D2 family homolog in hagfish. Our phylogenetic analysis showed that one of the two hagfish dopamine receptor D1 (Drd1×1) was placed in the basal position of D1A subfamily, while the other (Drd1×2) forms a cluster with D1C subfamily along with lamprey Drd1 genes (Fig. 2F). The hagfish D2 family homolog was in the Drd4 subfamily (thus, we classified it as a hagfish Drd4), suggesting that hagfish lacks Drd2. We next checked microsynteny conservation in representative vertebrate species. As a result, we found a considerably conserved pattern among the lamprey and jawed vertebrates, but the hagfish genome did not show any clear synteny blocks for the homologous genes (Suppl. Fig. 2). We thus suppose hagfish lacks Drd2, although our results did not directly demonstrate it.

### Molecular profiling of hagfish brain regions

Based on the results from our gene search and phylogenetic analysis, we decided to analyze gene expression patterns of *VGluT*, *GAD1*, *SerT*, *ChAT*, *TH*, *DATx1*, *DATx2*, *Drd1×1*, *Drd1×2*, and *Drd4* in the hagfish brain by *in situ* hybridization of cross-sections. In addition, we conducted AChE histochemistry to visualize both cholinergic neuronal somata and fibers. This technique chemically detects intrinsic AChE activity by *in situ* precipitation of copper ferrocyanide (Karnovsky & Roots, 1964). For this molecular profiling, we made a thorough series of transverse sections from the olfactory bulb to the rhombencephalon, as shown in Fig. 1C. The gene expression patterns in each brain region are summarized in Table 2.

**Table 2.**
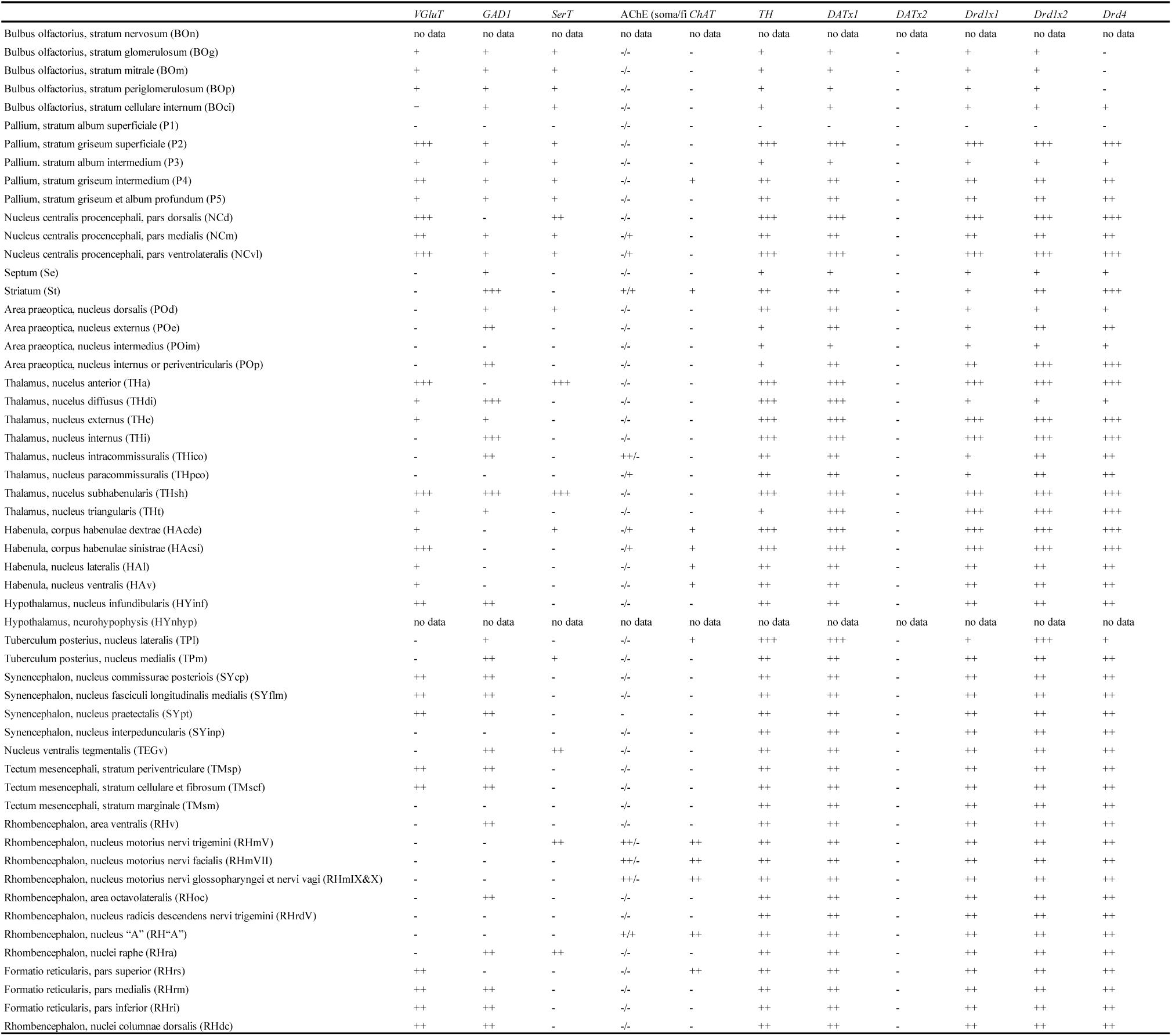
Gene expression patterns in each brain region.

#### Olfactory bulb

According to Wicht & Northcutt (1992), the burbus olfactorius (olfactory bulb, BO) is divided into five regions: stratum cellulare internum (BOci), glomerulosum (BOg), mitrale (BOm), nervosum (BOn), and periglomerulosum (BOp). Among them, BOn consists of a thin layer of olfactory nerve fibers, which is easily lost during sample collection, and thus was omitted from our analysis.

Notably, *VGluT* and *GAD1* showed distinct expression patterns; the former was expressed chiefly in the BOp (Fig. 3B), while the latter expression was predominantly observed in the innermost layers (i.e., the BOm and BOci; Fig. 3–4C). We also found that dopamine markers (*DATx1* and *TH*) and D1 receptor genes (*Drd1×1* and *Drd1×2*) were globally expressed in the BO (Fig. 3G, H, J, K), although not conspicuous in each glomerulus (only a few neurons are found in this structure, as suggested by Nissl staining; Fig. 3A). A similar expression was found for SerT (Fig. 3D). In contrast, no *ChAT*, *DATx2*, nor *Drd4* expression was found in this region (Fig. 3F, I, and L).

**Figure 3.**
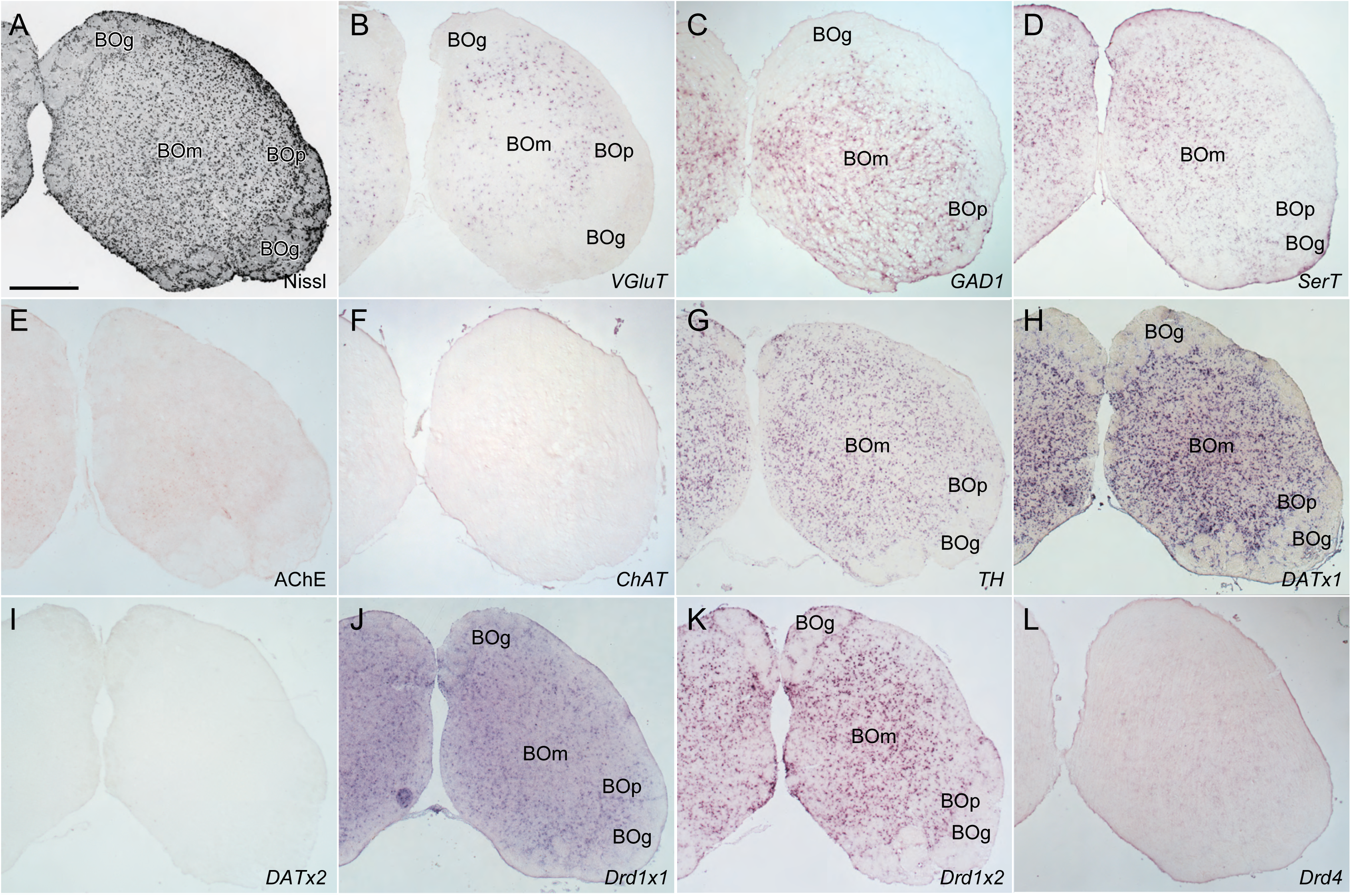
Histological and gene expression analysis in transversal sections at the anterior olfactory bulb. (A) Nissl staining, (B) *VGluT*, (C) *GAD1*, (D) *SerT*, (E) AChE staining, (F) *ChAT*, (G) *TH*, (H) *DATx1*, (I) *DATx2*, (J) *Drd1×1*, (K) *Drd1×2*, (L) *Drd4*. Scale bar: 500 μm in (A) for (A–L).

**Figure 4.**
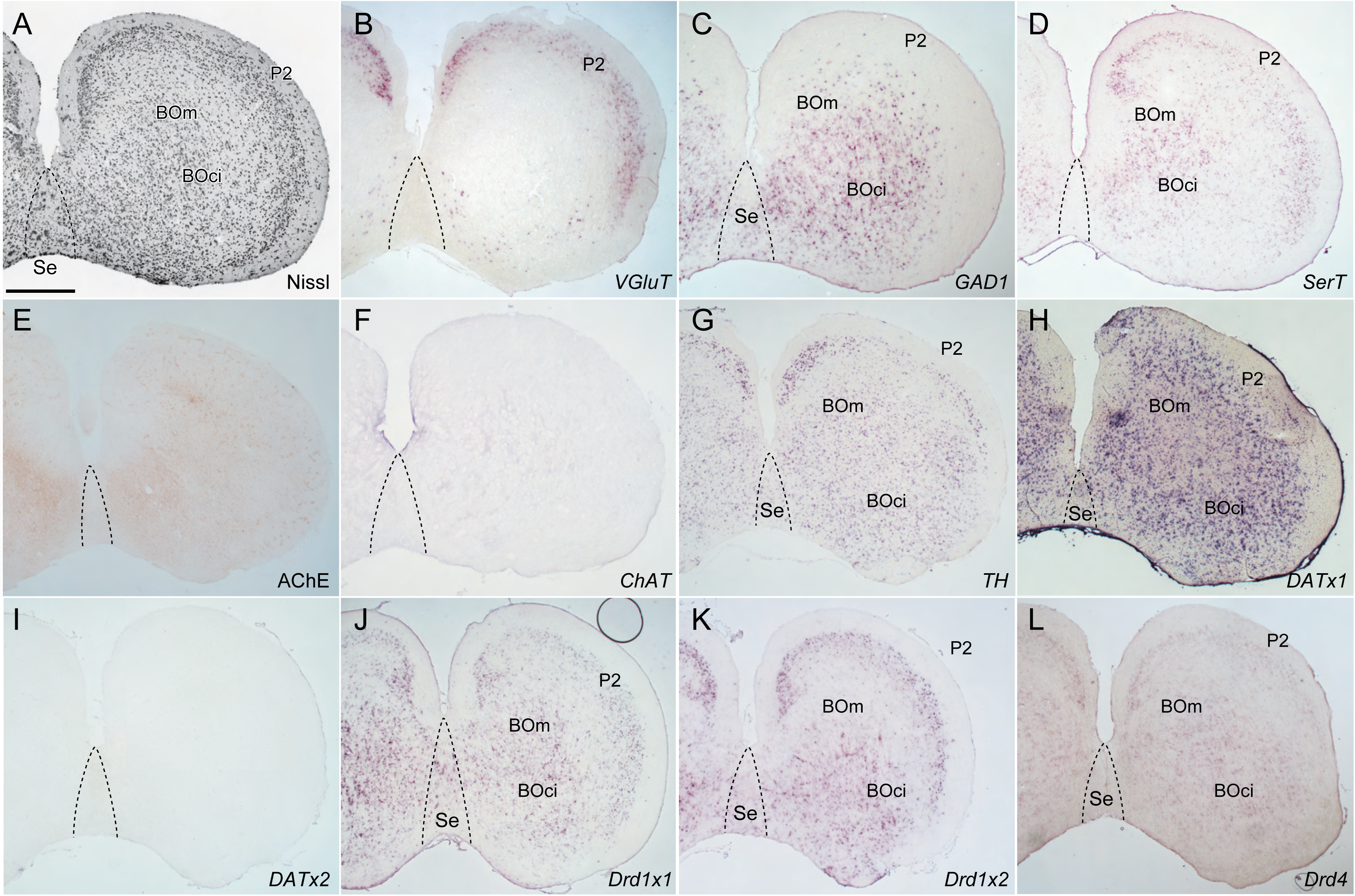
Histological and gene expression analysis in transversal sections at the posterior olfactory bulb. (A) Nissl staining, (B) *VGluT*, (C) *GAD1*, (D) *SerT*, (E) AChE staining, (F) *ChAT*, (G) *TH*, (H) *DATx1*, (I) *DATx2*, (J) *Drd1×1*, (K) *Drd1×2*, (L) *Drd4*. Scale bar: 500 μm in (A) for (A–L).

#### Telencephalon and diencephalon

In vertebrates, the telencephalon is generally divided into three major regions along the dorsoventral axis: the pallium, subpallium, and septum (Wicht & Northcutt, 1992).

In hagfish septum, no *VGluT* expression was observed. In contrast, *GAD1* positive cells were distributed in this region, especially in its medial part (Fig. 4C). In addition, the expression of *DATx1*, *Drd1×1*, *Drd1×2*, and *Drd4* was also observed (Fig. 4H, J, K). *TH* was also expressed sparsely in the septum, restricted in the lateral part (Fig. 4G).

The hagfish pallium consists of five layers named P1–P5, arranged from superficial to deep (Wicht & Northcutt, 1992). In the most superficial and predominantly fibrous P1 layer, no expression was detected for any genes we examined (Fig. 5–9). In the pallium, *VGluT* was strongly expressed in P2 and P4 (Fig. 4–8B), while GAD1 was expressed sparsely in P2 to P5 (Fig. 4–8C). *SerT* was expressed mainly in P2 (Fig. 4–9D). *TH*, *DATx1, Drd1×1, Drd1×2,* and *Drd4* expressions were observed in P2–P5 but especially stronger in P2 and weaker in P3, possibly because of the difference of cell density. AChE-positive fibers were observed in the commusura interbulbaris (coib; Fig 7E, 8E), which connects the pallia of both hemispheres (Wicht & Nieuwenhuys, 1998).

**Figure 5.**
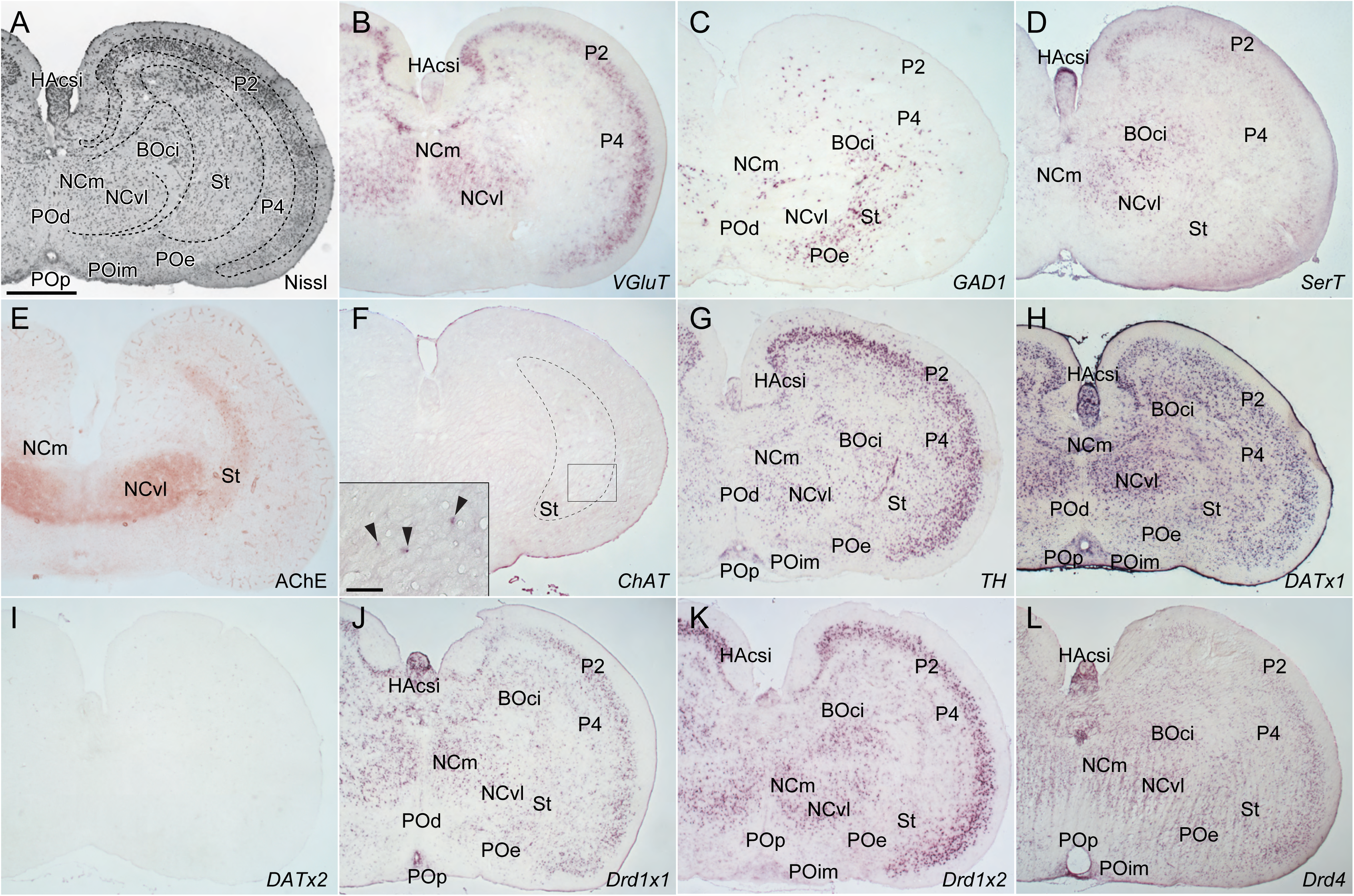
Histological and gene expression analysis in transversal sections at the anteriormost telencephalon. (A) Nissl staining, (B) *VGluT*, (C) *GAD1*, (D) *SerT*, (E) AChE staining, (F) *ChAT*, (G) *TH*, (H) *DATx1*, (I) *DATx2*, (J) *Drd1×1*, (K) *Drd1×2*, (L) *Drd4*. Scale bar: 500 μm in (A) for (A–L), 100 μm in inset (F).

**Figure 6.**
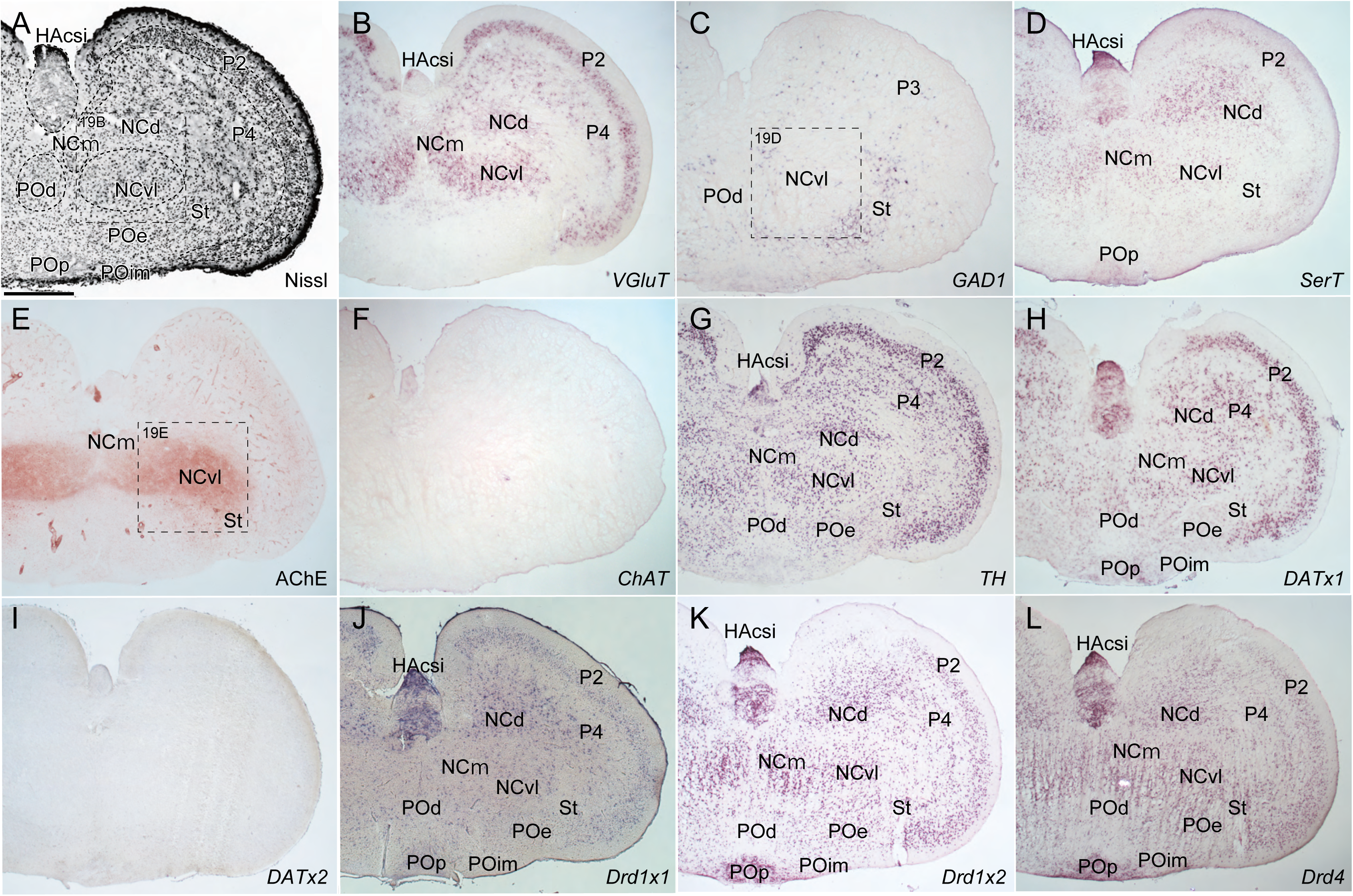
Histological and gene expression analysis in transversal sections at the mid-anterior telencephalon. (A) Nissl staining, (B) *VGluT*, (C) *GAD1*, (D) *SerT*, (E) AChE staining, (F) *ChAT*, (G) *TH*, (H) *DATx1*, (I) *DATx2*, (J) *Drd1×1*, (K) *Drd1×2*, (L) *Drd4*. Scale bar: 500 μm in (A) for (A–L).

**Figure 7.**
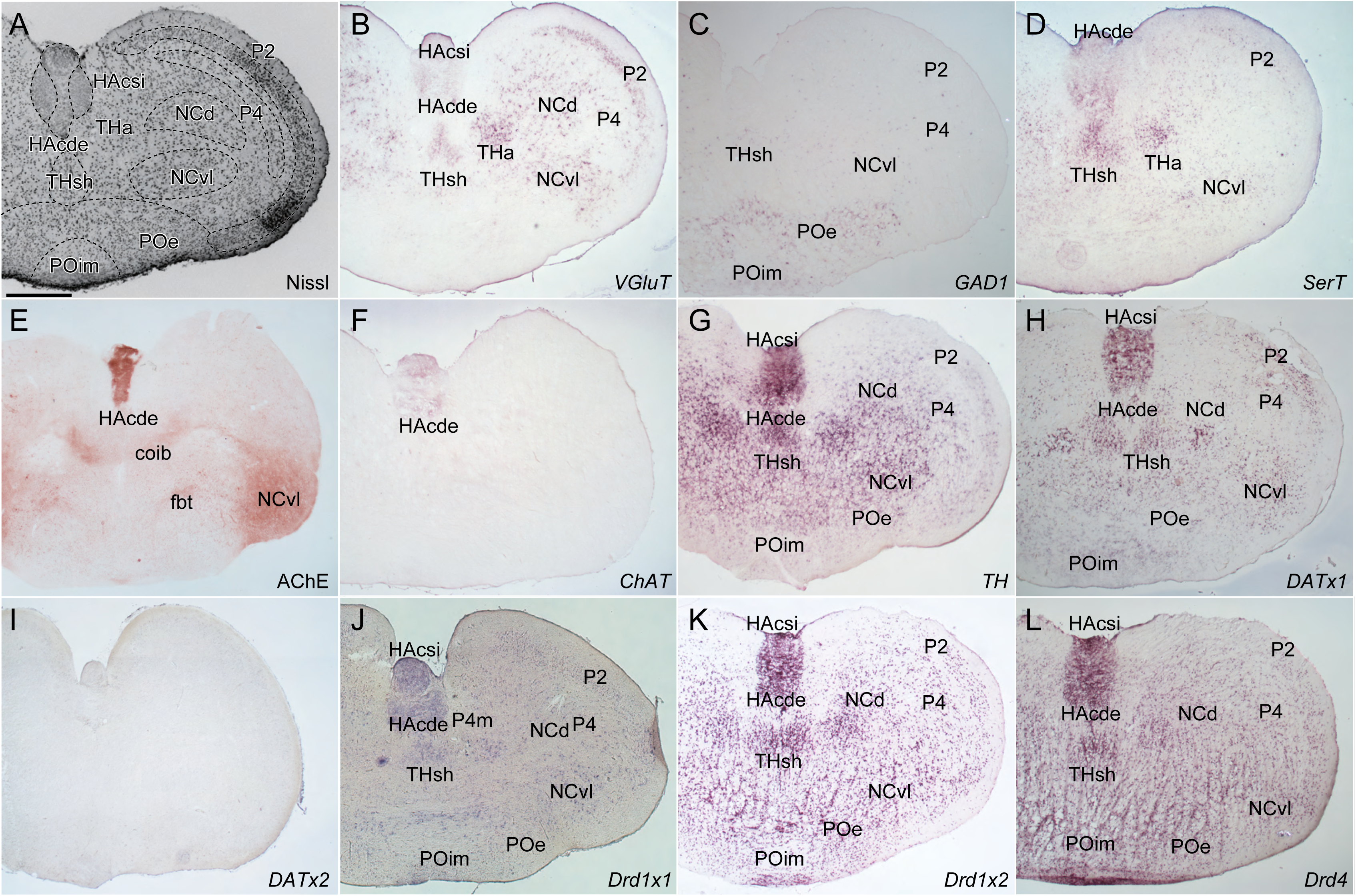
Histological and gene expression analysis in transversal sections at the mid-posterior telencephalon. (A) Nissl staining, (B) *VGluT*, (C) *GAD1*, (D) *SerT*, (E) AChE staining, (F) *ChAT*, (G) *TH*, (H) *DATx1*, (I) *DATx2*, (J) *Drd1×1*, (K) *Drd1×2*, (L) *Drd4*. Scale bar: 500 μm in (A) for (A–L).

**Figure 8.**
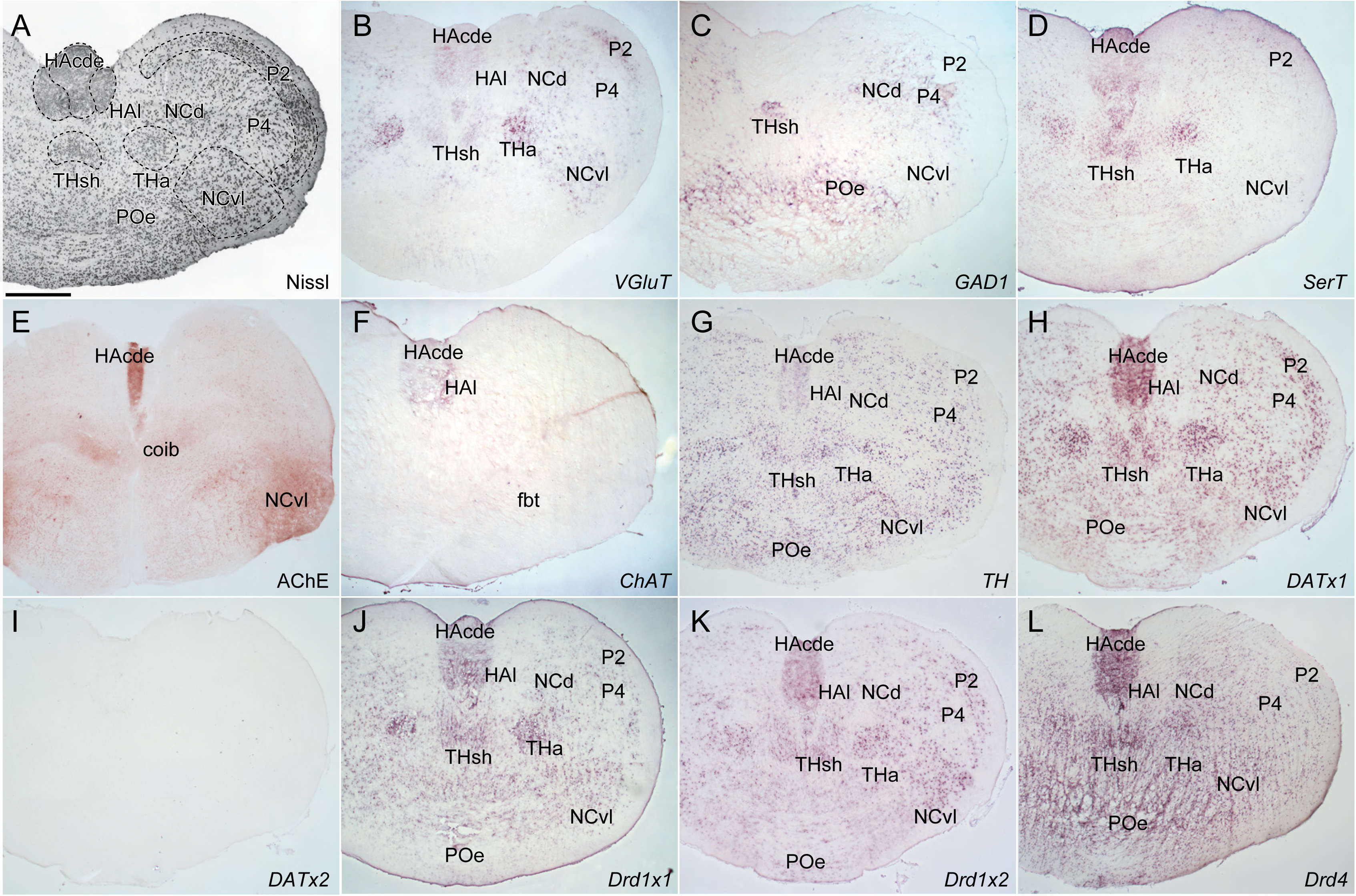
Histological and gene expression analysis in transversal sections at the posteriormost telencephalon. (A) Nissl staining, (B) *VGluT*, (C) *GAD1*, (D) *SerT*, (E) AChE staining, (F) *ChAT*, (G) *TH*, (H) *DATx1*, (I) *DATx2*, (J) *Drd1×1*, (K) *Drd1×2*, (L) *Drd4*. Scale bar: 500 μm in (A) for (A–L).

**Figure 9.**
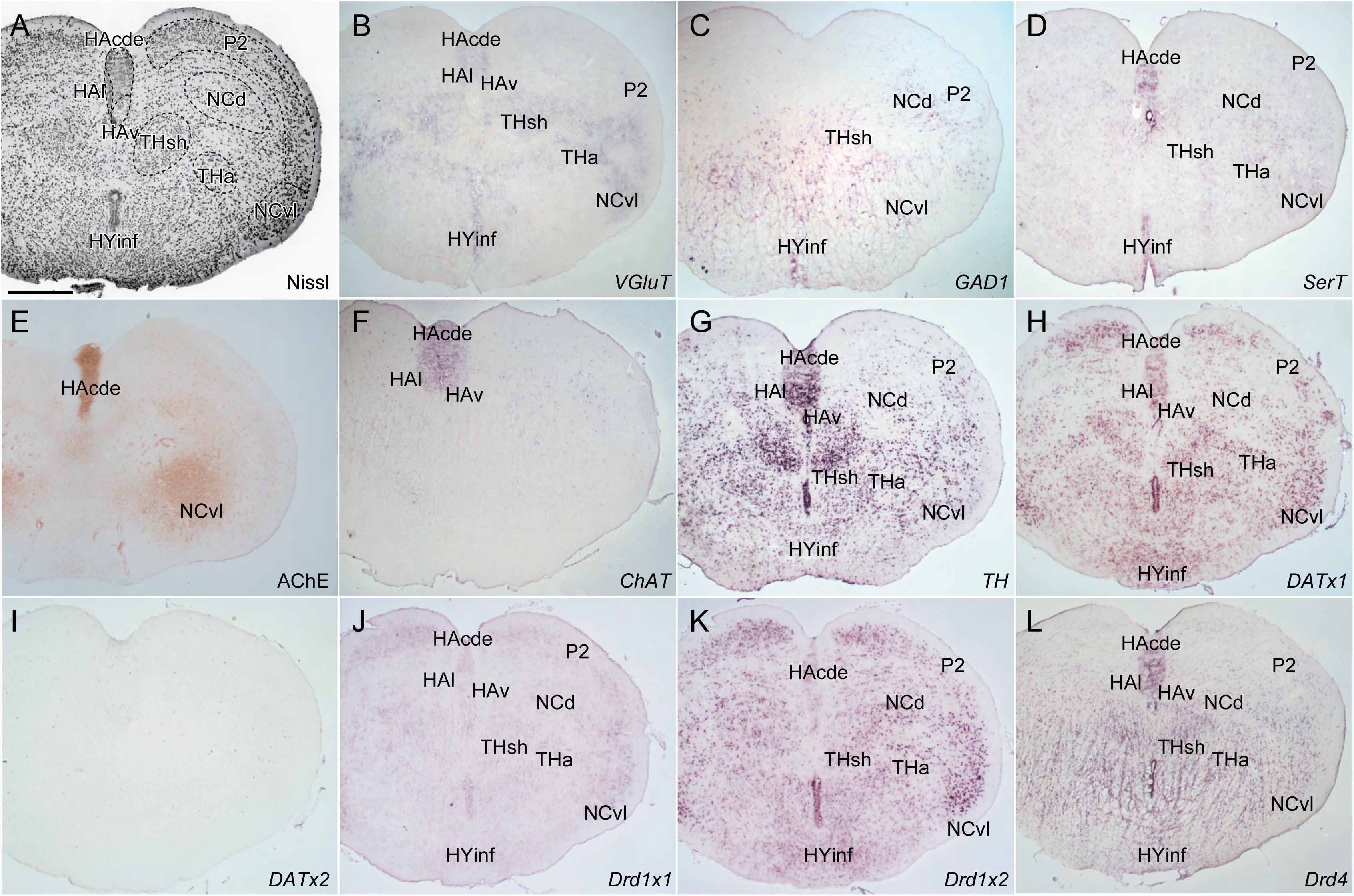
Histological and gene expression analysis in transversal sections at the anteriormost diencephalon.; (A) Nissl staining, (B) *VGluT*, (C) *GAD1*, (D) *SerT*, (E) AChE staining, (F) *ChAT*, (G) *TH*, (H) *DATx1*, (I) *DATx2*, (J) *Drd1×1*, (K) *Drd1×2*, (L) *Drd4*. Scale bar: 500 μm in (A) for (A–L).

In the striatum identified by Wicht & Northcutt (1992), no *VGluT* expression was observe (Fig. 5–6B) but *GAD1* was strongly expressed (Fig. 5–6C). Some *SerT*-expressing cells were observed in this brain region (Fig. 5–6D). Both AChE positive cell bodies and fibers were found in the striatum (Fig. 5E). Consistently, sparse *ChAT* expression was observed in this region (Fig. 5F). All dopamine-related genes except for *DATx2* were also expressed in the striatum (Fig. 5–6, G–L).

The brain region called “nucleus centralis procencephali” or the central prosencephalic nucleus is an idiosyncratic structure in the hagfish brain (Wicht & Nieuwenhuys, 1998). Wicht & Northcutt (1992) subdivided this structure into pars medialis (NCm), ventrolateralis (NCvl), and dorsalis (NCd). *VGluT* was uniformly expressed in all these subnuclei except for the posteriormost part of NCd (Fig. 6–8B), while different expression patterns in each subnucleus were found for *GAD1*; no expression in anterior NCd, densely expressed in posterior NCd and NCm, and sparsely expressed in NCvl (Fig. 6–8C). *SerT* was expressed in NCm, NCvl, and NCd (Fig. 6–9D). We observed AChE-positive fibers in NCm and NCvl (albeit stronger in the latter; Fig. 5−6E) as well as in the fasculus basalis telencephali (fbt; Fig. 7E). All dopamine-related genes except for *DATx2* were also expressed in all subnuclei (Fig. 5–6, G–L).

According to Wicht & Northcutt (1992), the area praeoptica (the preoptic area, PO) of hagfish consists of four nuclei, the nucleus externus (POe), dorsalis (POd), intermedius (POim), and internus or periventricularis (POp). No *VGluT* expression was found in this area (Fig. 5–8B), and *GAD1* was expressed in the POe and POd (Fig. 5–8C). We observed *SerT* expression in Pop (Fig. 6D). As for dopamine-related genes, *TH* was sparsely expressed in the entire PO (Fig. 5–8G). In contrast, we observed dense *DATx1* expression in all four nuclei (Fig. 5–8H). No evident *DATx2* expression was found (Fig. 5–8I). *Drd1×1* expression was weak in general but stronger in POp (Fig. 5–8J). We observed *Drd1×2* and *Drd4* expression in all four nuclei, relatively weak in the POd and significantly strong in POp (Fig. 5–8, K and L).

Wicht & Northcutt (1992) differentiates the hagfish thalamus into eight nuclei; the nucleus anterior (THa; possibly corresponds to the lateral genuculate nucleus, LGN, of the amniotes because this region receives retinal inputs; Wicht & Northcutt, 1990), diffusus (THdi), externus (THe), internus (THi), intracommisularis (THico), paracommisularis (THpco), subhabenularis (THsh), triangularis (THt). While *VGluT* was strongly expressed in the THa (Fig. 8–9B), *GAD1* showed strong expression in the THdi and THi (Fig. 9–12C). Both genes are expressed in the THsh and THt in a similar manner (Fig. 8–12, B, C). Notably, we observed *SerT* expression in the THa and THsh (Fig. 8–9D). Both AChE-positive cell bodies and fibers were found in the THpco (Fig.10E), consistent with sparse expression of *ChAT* (Fig.10F), possibly corresponding to the ventromedial and ventrolateral nuclei of the fish thalamus (Clemente et al., 2004). All dopamine-related genes except for *DATx2* were broadly expressed in the entire thalamus (Fig. 7–9, G–L).

**Figure 10.**
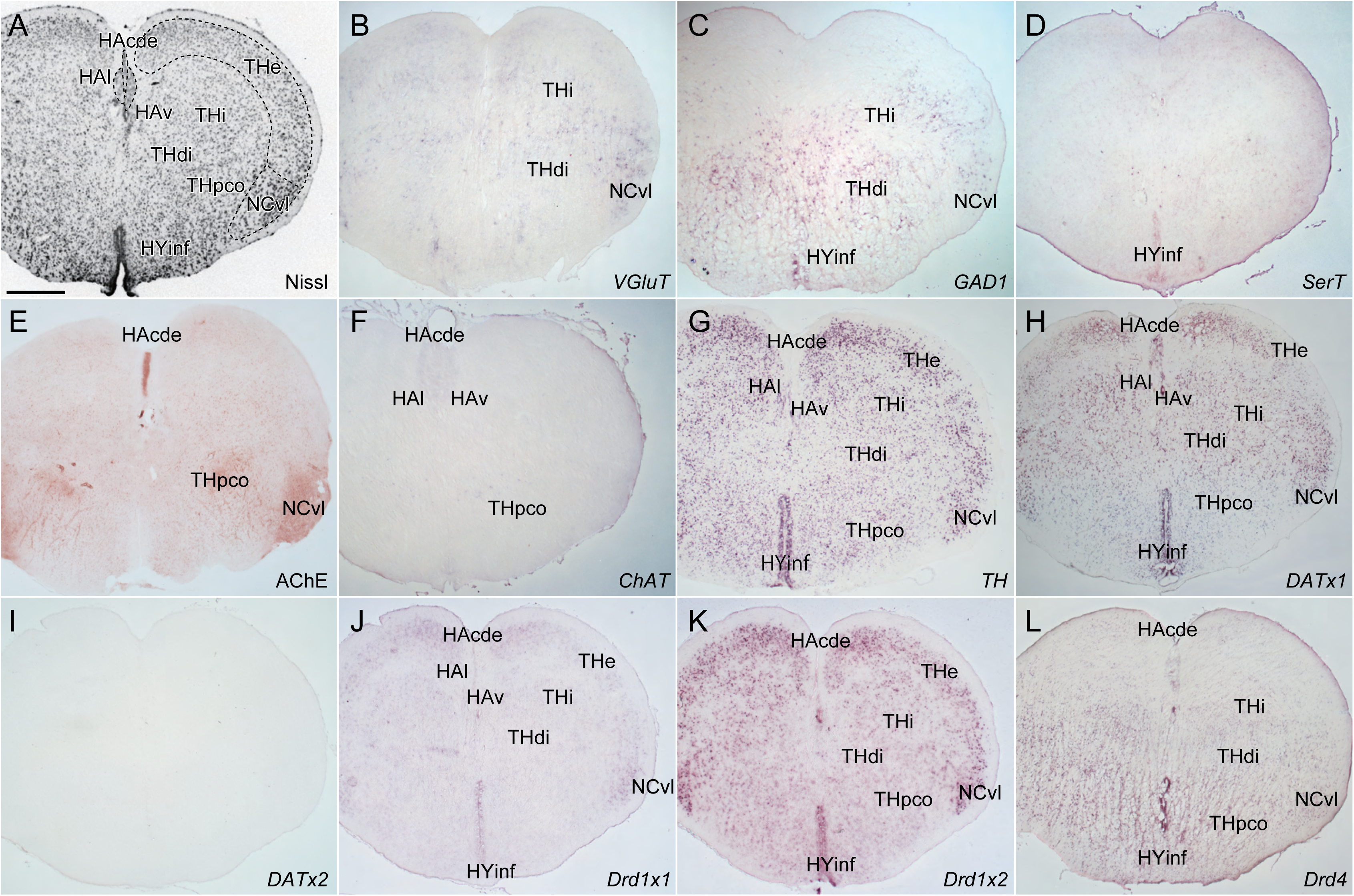
Histological and gene expression analysis in transversal sections at the mid-anterior diencephalon. (A) Nissl staining, (B) *VGluT*, (C) *GAD1*, (D) *SerT*, (E) AChE staining, (F) *ChAT*, (G) *TH*, (H) *DATx1*, (I) *DATx2*, (J) *Drd1×1*, (K) *Drd1×2*, (L) *Drd4*. Scale bar: 500 μm in (A) for (A–L).

**Figure 11.**
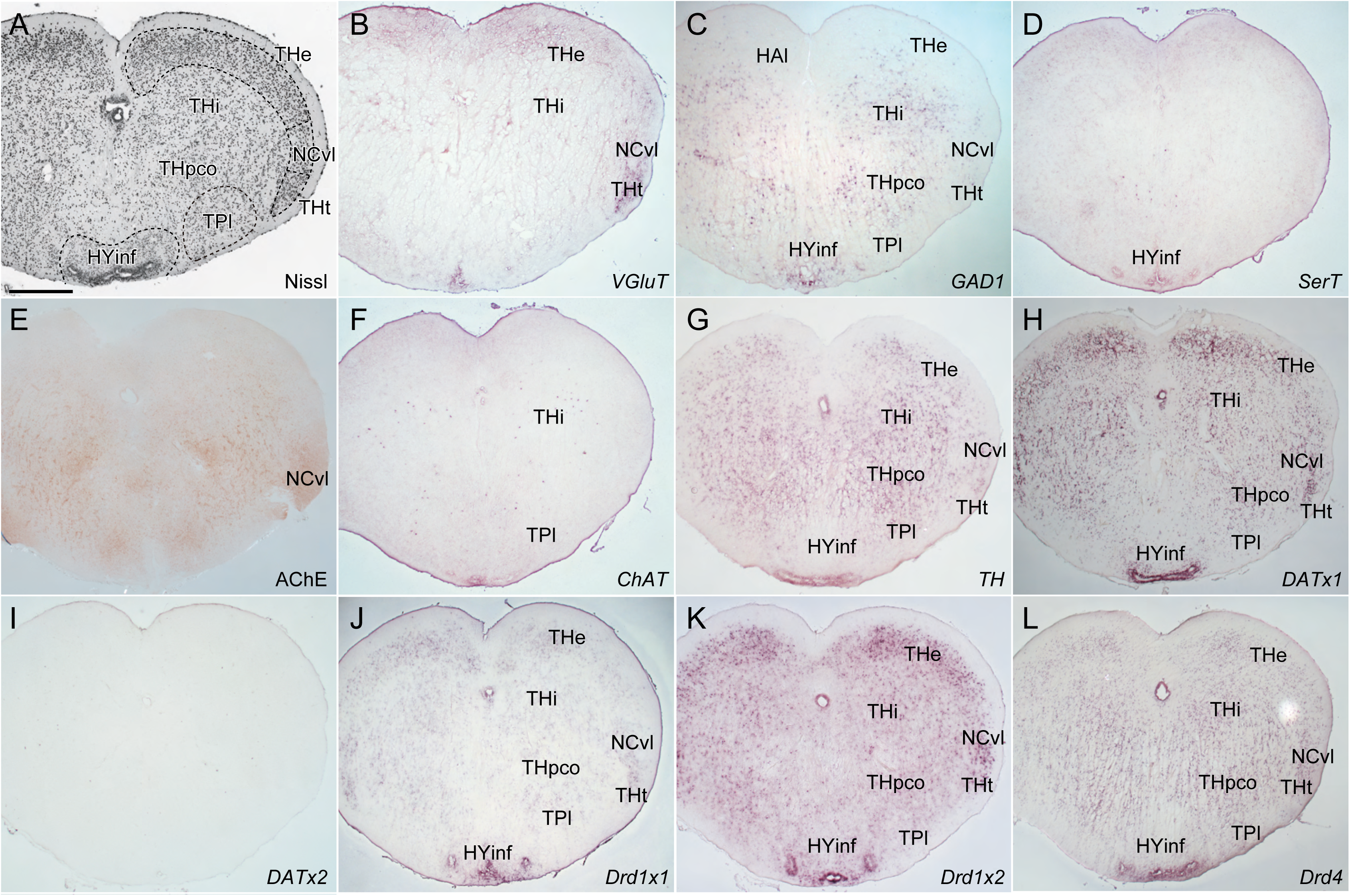
Histological and gene expression analysis in transversal sections at the mid-posterior diencephalon. (A) Nissl staining, (B) *VGluT*, (C) *GAD1*, (D) *SerT*, (E) AChE staining, (F) *ChAT*, (G) *TH*, (H) *DATx1*, (I) *DATx2*, (J) *Drd1×1*, (K) *Drd1×2*, (L) *Drd4*. Scale bar: 500 μm in (A) for (A–L).

**Figure 12.**
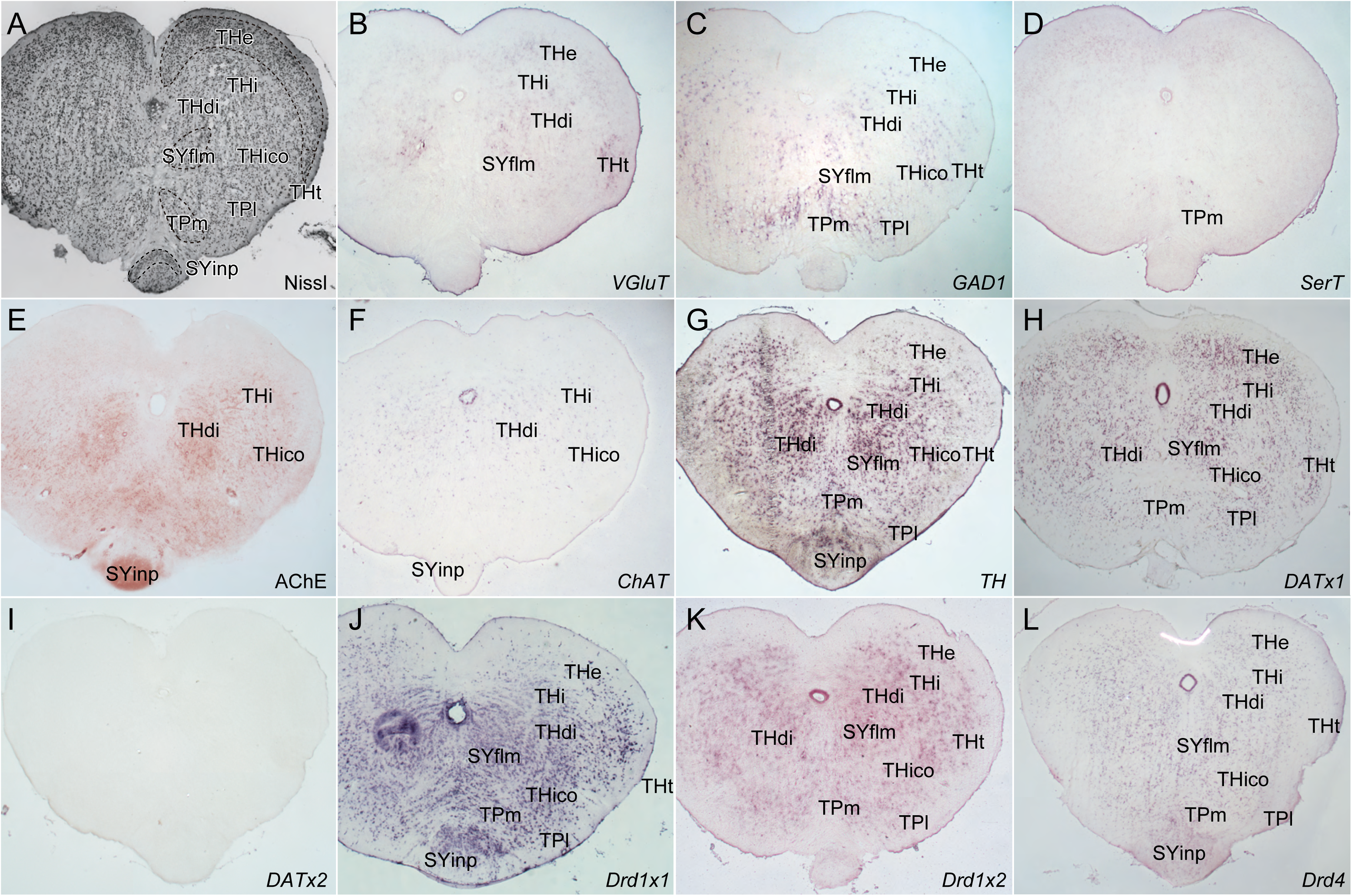
Histological and gene expression analysis in transversal sections at the posteriormost diencephalon. (A) Nissl staining, (B) *VGluT*, (C) *GAD1*, (D) *SerT*, (E) AChE staining, (F) *ChAT*, (G) *TH*, (H) *DATx1*, (I) *DATx2*, (J) *Drd1×1*, (K) *Drd1×2*, (L) *Drd4*. Scale bar: 500 μm in (A) for (A–L).

The hagfish habenula (HA) can be subdivided into four parts (Wicht & Northcutt, 1992); the corpus habenulae dextrae (HAcde), corpus habenulae sinistrae (HAcsi), nucleus latelaris (HAl), and nucleus ventralis (HAv). We observed *VGluT* expression in the entire habenula (Fig. 7–9B), although no *GAD1* expression was found in the habenula (Fig. 7–9C). The expression of *SerT* was observed only in the HAcde, while *ChAT* was expressed in the entire habenula (Fig7–9, D, F). AChE-positive fibers were found in the medial part of the HAcde (Fig7–10E). All dopamine-related genes except for *DATx2* were broadly expressed in the entire habenula (Fig7–9, G–L).

Wicht & Northcutt (1992) subdivided the hagfish hypothalamus (HY) into the nucleus infundibularis (HYinf) and the nucleus neurohypophysis (HYnhyp). However, the latter structure was not observed during our preparations because it is embedded in the extraencephalic tissue (Yamaguchi et al., 2023). In the HYinf, *VGluT* was expressed only in its periventricular cells, while the expression of *GAD1* was observed in both periventricular and non-periventricular part (Fig. 9–11B, C). In addition, *SerT* was observed to be expressed only in the periventricular cells (Fig. 9–11D). Among the dopamine-related genes, *TH*, *DATx1*, *Drd1×2*, *Drd4* were expressed both in the periventricular and non-periventricular cells (Fig. 9–11, G–H, K–L) but *Drd1×1* expression was observed only in the periventricular cells (Fig. 9–11J). We found no *DATx2* expression in this region (Fig. 9–11I).

The tuberculum posterius (the posterior tuberculum, TP) is subdivided into two regions (Wicht & Northcutt, 1992; Wicht & Nieuwenhuys, 1998); the nucleus lateralis (TPl) and medialis (TPm). No *VGluT* expression was observed, but *GAD1* expression was confirmed in these regions (Fig. 11–12B, C). *ChAT* was scatteredly expressed in the TPl (Fig. 11–12F), while the expression of *SerT* was found in the TPm (Fig. 11–12D). Except for *DATx2*, All dopamine-related genes except for *DATx2* were expressed in both TPl and TPm (Fig. 11–12, G–L).

The synencephalon is the caudal-most part of the diencephalon and corresponds to the embryonic prosomere 1 (Ferran et al., 2007; Lauter et al., 2013). Wicht & Northcutt (1992) assume that the hagfish synencephalon is composed of the nucleus fasciculi longitudinalis medialis (SYflm), the nucleus commissurae posteriois (SYcp), the nucleus praetectalis (SYpt), and the nucleus interpeduncularis (SYinp). Both *VGluT* and *GAD1* were expressed in SYflm, SYcp, and SYpt (Fig. 12–13, B, C), as well as all dopamine-related genes except for *DATx2* (Fig. 12–13, G–L). AChE-positive somata and fibers were observed in SYinp, where *ChAT* were also scatteredly expressed (Fig. 12–13E, 12F). In the SYinp, no expression was observed for either *VGluT* or *GAD1* (Fig. 12–13, B–C), while all dopamine-related genes except for *DATx2* were expressed (Fig. 12–13, G–L). AChE-positive fibers were observed here.

**Figure 13.**
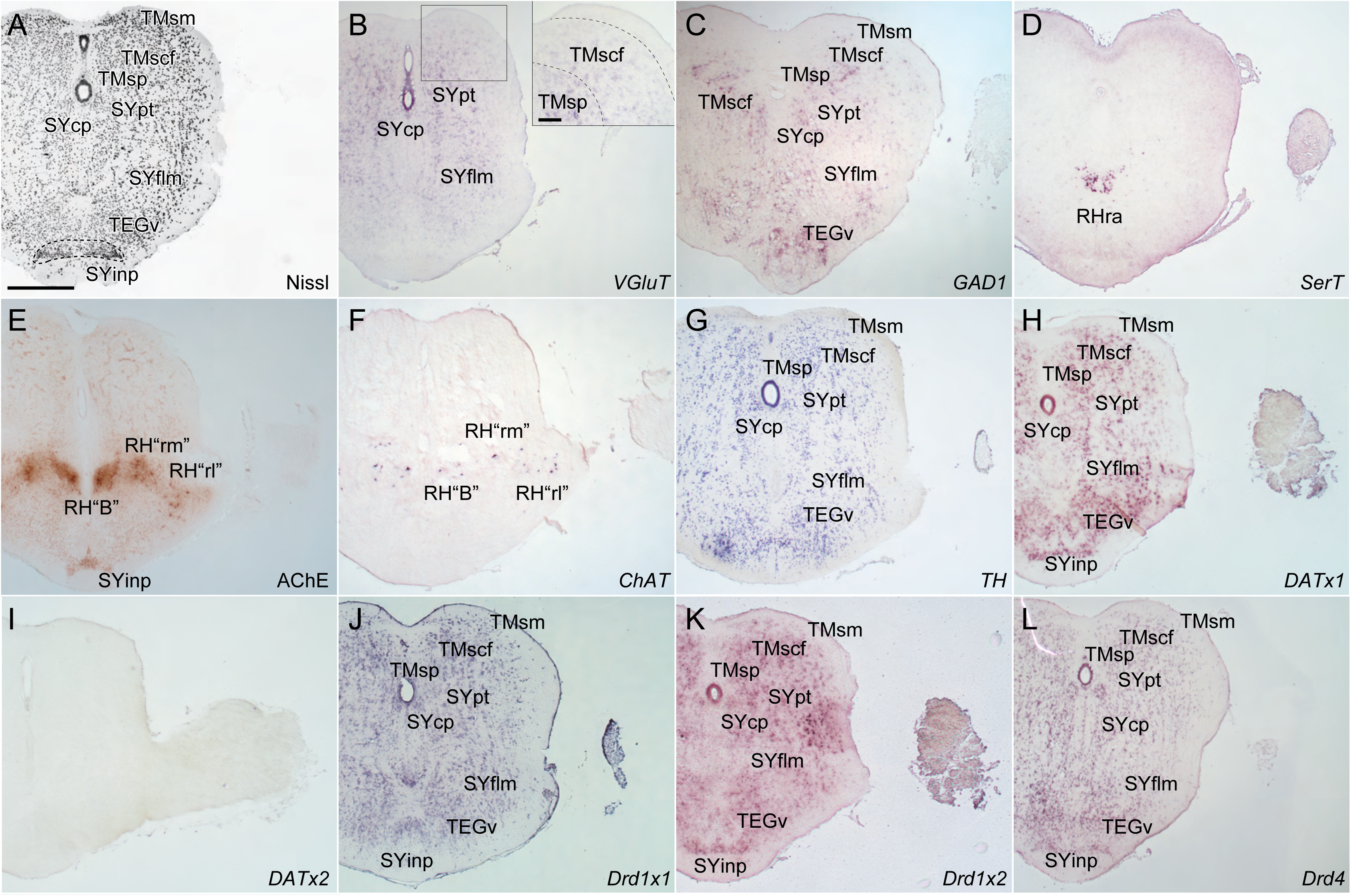
Histological and gene expression analysis in transversal sections at the anterior mesencephalon. (A) Nissl staining, (B) *VGluT*, (C) *GAD1*, (D) *SerT*, (E) AChE staining, (F) *ChAT*, (G) *TH*, (H) *DATx1*, (I) *DATx2*, (J) *Drd1×1*, (K) *Drd1×2*, (L) *Drd4*. Scale bar: 500 μm in (A) for (A–L), 100 μm in inset (F).

#### Mesencephalon

The tectum mesencephali (the mesencephalic tectum, TM) is composed of three major layers (Iwahori et al., 1996); the stratum periventriculare (TMsp), the stratum cellulare et fibrosum (TMscf), and the stratum marginale (TMsm). Among these layers, both *VGluT* and *GAD1* expression was found in the TMsp and TMscf (Fig. 13–14, B–C). Furthermore, all dopamine-related genes except for *DATx2* were also expressed in these three layers (Fig. 13–14, G–L).

**Figure 14.**
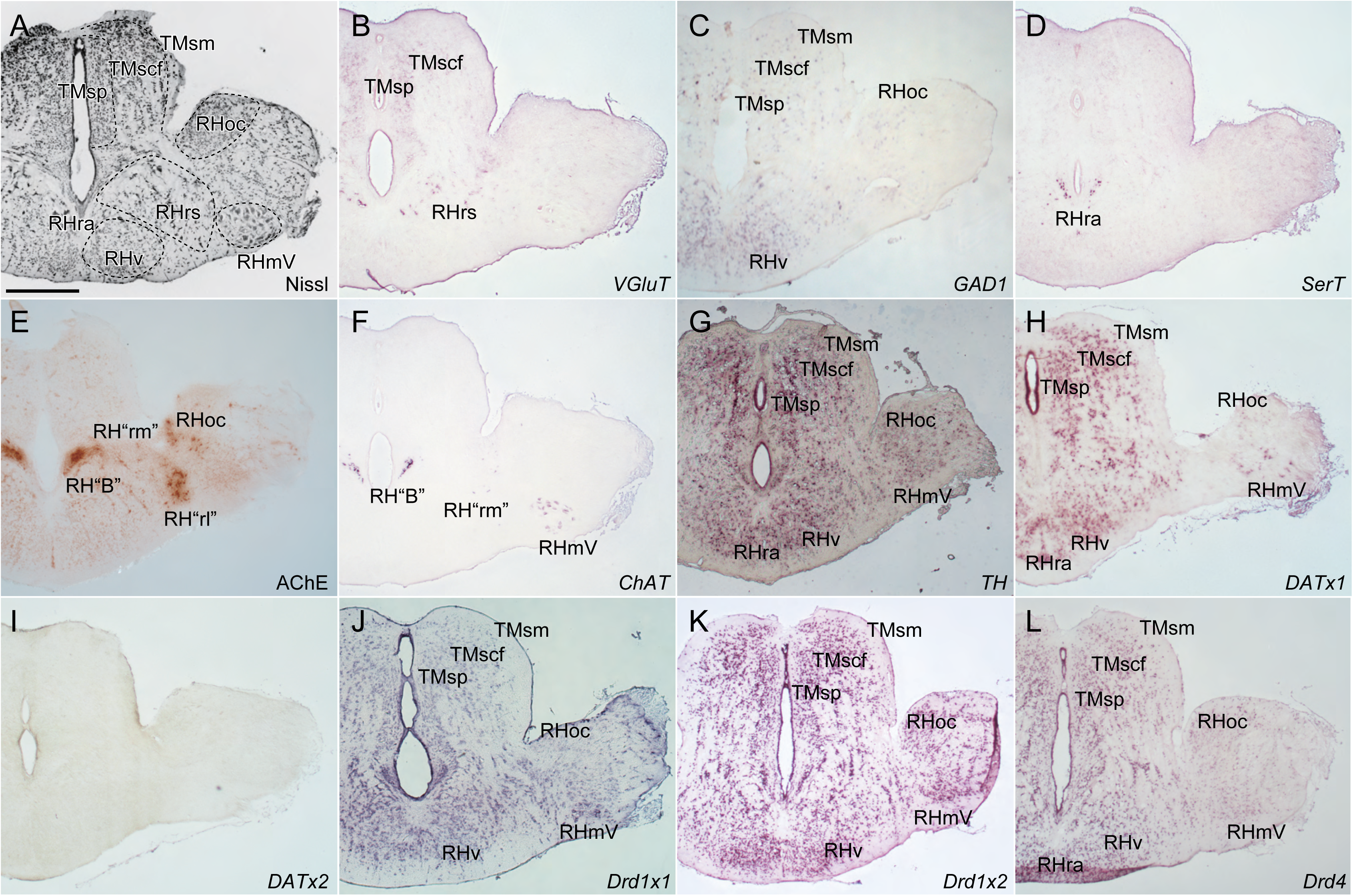
Histological and gene expression analysis in transversal sections at the mid-mesencephalon. (A) Nissl staining, (B) *VGluT*, (C) *GAD1*, (D) *SerT*, (E) AChE staining, (F) *ChAT*, (G) *TH*, (H) *DATx1*, (I) *DATx2*, (J) *Drd1×1*, (K) *Drd1×2*, (L) *Drd4*. Scale bar: 500 μm in (A) for (A–L).

In the midbrain tegmentum (TEG), Wicht & Northcutt (1992) described only one nucleus: the nucleus ventralis tegmentalis (TEGv). No *VGluT* but *GAD1* expression was observed in this nucleus (Fig. 13B, C). All dopamine-related genes except for *DATx2* were also expressed (Fig. 13G–L).

#### Rhombencephalon

For the hindbrain reticular formation, different nomenclatures are used in Wicht & Northcutt (1992) and Wicht & Nieuwenhuys (1998). In particular, Wicht & Northcutt (1992) named one region as nuclei reticulares (Ret), while Wicht & Nieuwenhuys (1998) as tegmentum ventrale (tegmv). In the present study, we labeled it as the ventral area (RHv) of the rhombencephalon. We observed no *VGluT* but *GAD1* expression in this region (Fig. 13–15, B, C). All dopamine-related genes except for *DATx2* were also expressed here.

**Figure 15.**
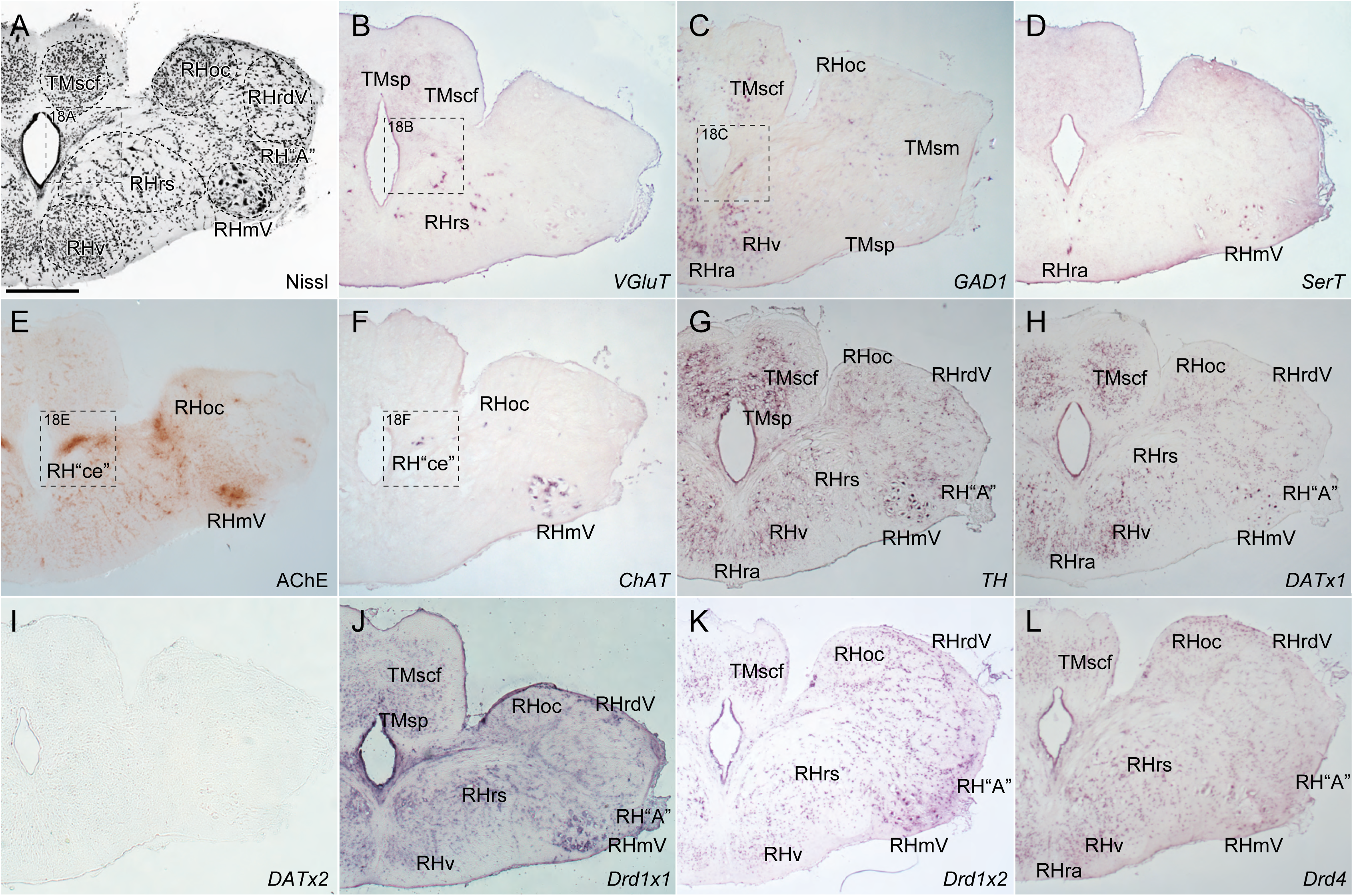
Histological and gene expression analysis in transversal sections at the posterior mesencephalon/anterior rhombencephalon. (A) Nissl staining, (B) *VGluT*, (C) *GAD1*, (D) *SerT*, (E) AChE staining, (F) *ChAT*, (G) *TH*, (H) *DATx1*, (I) *DATx2*, (J) *Drd1×1*, (K) *Drd1×2*, (L) *Drd4*. Scale bar: 500 μm in (A) for (A–L).

Apart from the RHv, we basically followed the nomenclature in Wicht & Nieuwenhuys (1998) hereafter, because it provides a more detailed description of hindbrain regions than Wicht & Northcutt (1992). For the comparison of the nomenclature, see Suppl. Table. 1.

In nucleus radicis descendens nervi trigemini (RHrdV), *VGluT*- and *GAD1*-positive cells were found at its caudal-most level in a scattered manner (Fig. 15–17, B, C). In contrast, all dopamine-related genes except for *DATx2* were expressed throughout this region (Fig. 15–17, G–L).

**Figure 16.**
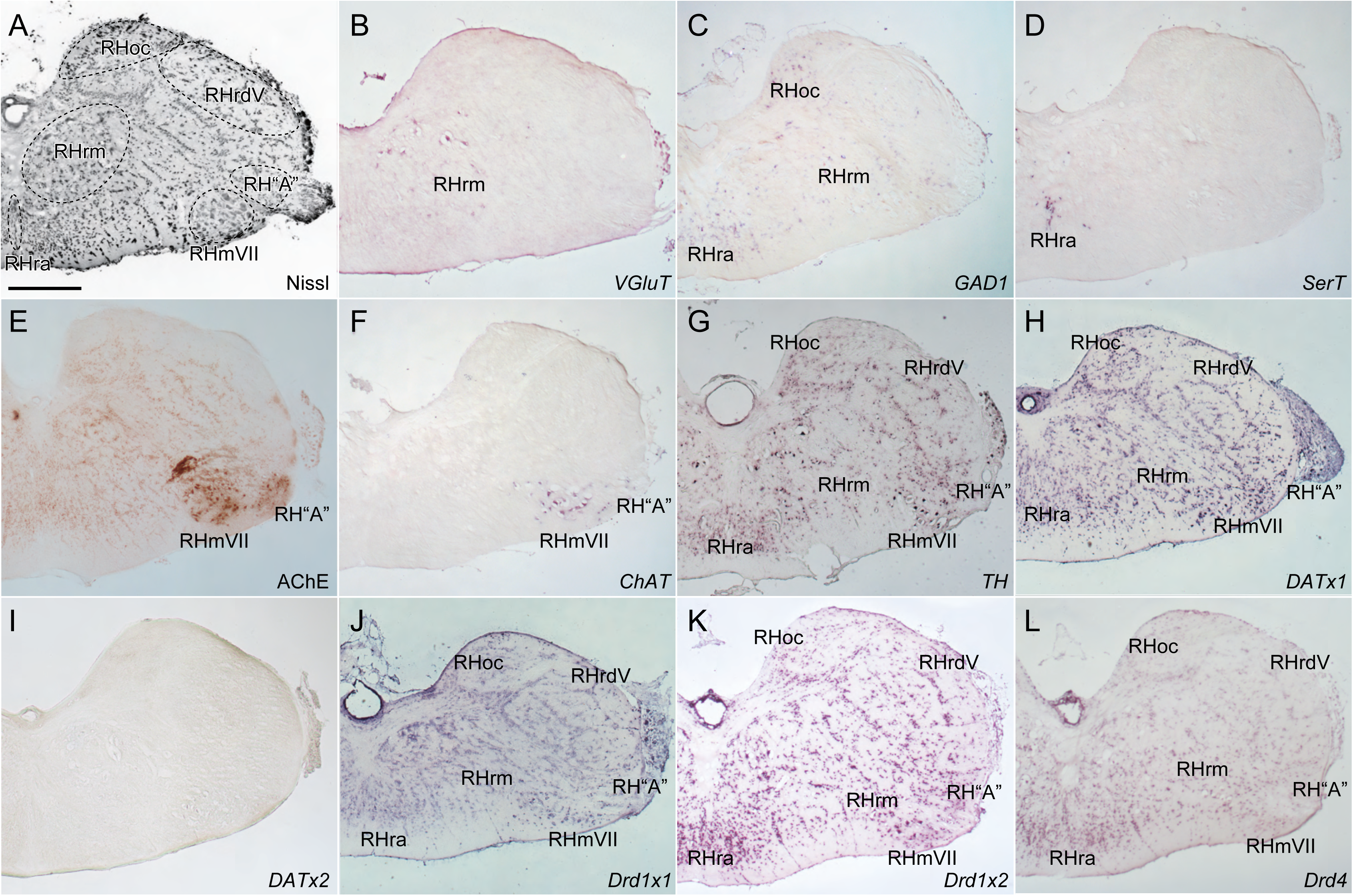
Histological and gene expression analysis in transversal sections at the mid-rhombencephalon. (A) Nissl staining, (B) *VGluT*, (C) *GAD1*, (D) *SerT*, (E) AChE staining, (F) *ChAT*, (G) *TH*, (H) *DATx1*, (I) *DATx2*, (J) *Drd1×1*, (K) *Drd1×2*, (L) *Drd4*. Scale bar: 500 μm in (A) for (A–L).

**Figure 17.**
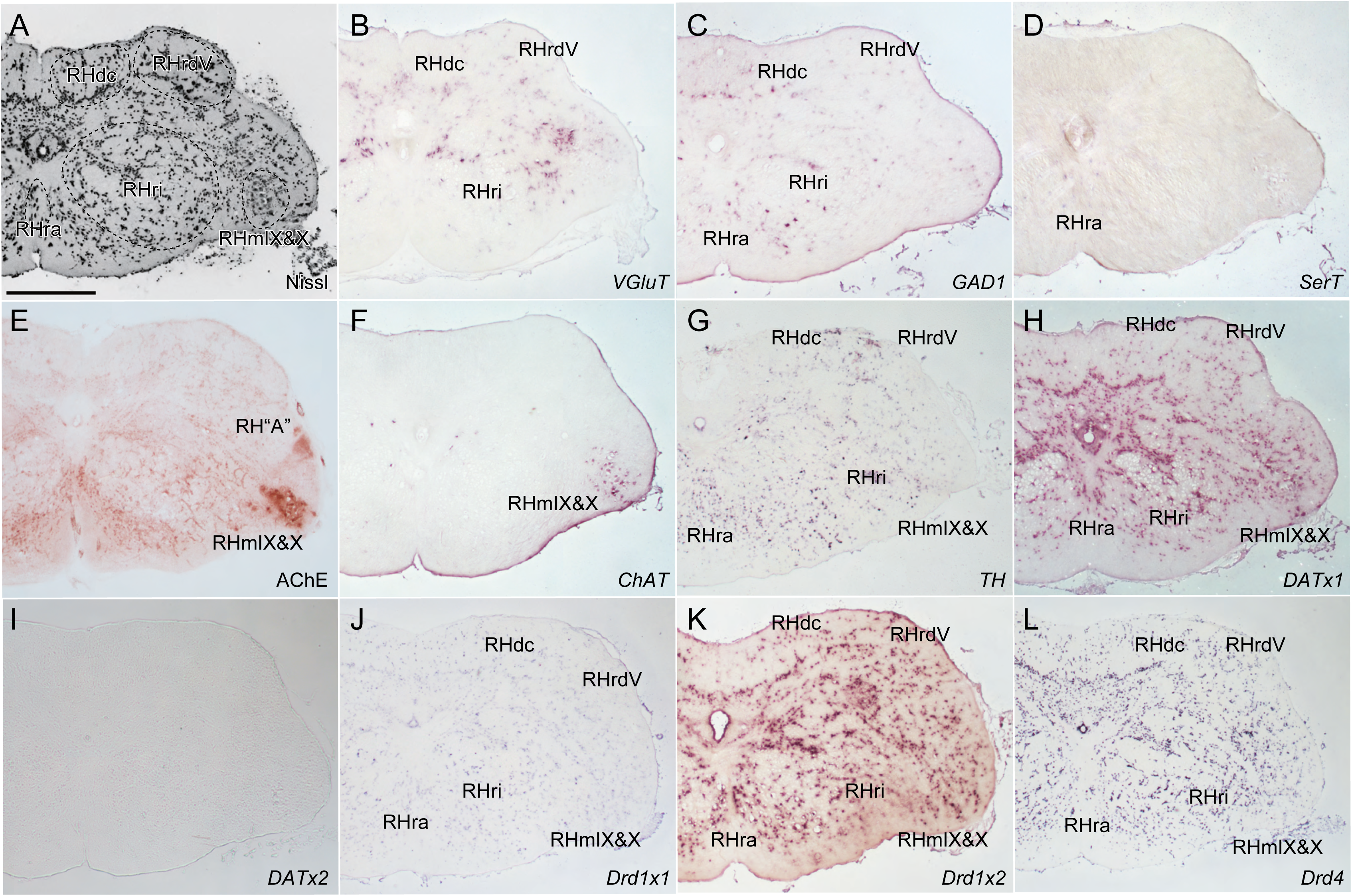
Histological and gene expression analysis in transversal sections at the posterior-rhombencephalon. (A) Nissl staining, (B) *VGluT*, (C) *GAD1*, (D) *SerT*, (E) AChE staining, (F) *ChAT*, (G) *TH*, (H) *DATx1*, (I) *DATx2*, (J) *Drd1×1*, (K) *Drd1×2*, (L) *Drd4*. Scale bar: 500 μm in (A) for (A–L).

In area octavolateralis (RHoc), no *VGluT* but *GAD1* expression was observed (Fig. 14–16, B, C). All dopamine dopamine-related genes except for *DATx2* were also expressed in this region (Fig. 14–16, G–L).

Wicht & Nieuwenhuys (1998) divided the hagfish reticular formation into pars superior (RHrs), pars medialis (RHrm), and pars inferior (RHri). *VGluT* was expressed in all these subregions, but *GAD1* expression was observed only in the RHrm and RHri (Fig. 14–17, B, C). Expression of all dopamine-related genes except for *DATx2* was consistently observed in these subregions (Fig. 14–17, G–L).

In the report of AChE staining by Kusunoki et al. (1982), they recognized several AChE-positive nuclei in the anterior rhonmencephalon: the rostromedialmedial/rostrolateral parts of the rhombencephalic reticular formation (RH“rm”/RH“rl”) and nucleus “B” (RH“B”). We could obtain the same results of AChE staining (Fig. 13–14E). Furthermore, we found ChAT-positive cells in these brain regions (Fig. 13–14E).

Kusunoki et al. (1982) also reported AChE staining signals in the primordial cerebellum (RH“ce”) described by Larsell (1967, 1947). We also found AChE-positive cells and fibers in the corresponding region, continuing from RH“B” (Fig. 15E). In this region, *ChAT*-positive cells were observed (Fig. 15F). We conducted further analysis for this region as described in the next section.

There is an enigmatic nucleus “A” of Kusunoki et al. (1982) in the hagfish rhombencephalon (RH“A”). Consistent with the result of Kusunoki et al. (1982), we confirmed AchE staining signals as well as *ChAT* expression in this nucleus (Fig. 14–15, E, F). No expression was observed for either *VGluT* or *GAD1* (Fig. 14–15, B, C). All dopamine-related genes except for *DATx2* were expressed here (Fig. 14–15, G–L).

In the nuclei raphe (RHra), a specific expression of *SerT* was found (Fig. 15–17D). In addition, no *VGluT* but *GAD1* expression was observed (Fig. 15–17, B, C). All dopamine dopamine-related genes except for *DATx2* were also expressed in this region (Fig. 15–17, G–L).

In the nuclei columnae dorsalis (RHdc), we observed gene expression for *VGluT* and *GAD1*, as well as all dopamine redopamine-related genes except for *DATx2* (Fig. 17, G–L).

Lastly, we confirmed dense signals of AchE staining and *ChAT* expression in the motor nuclei (nucleus motorius nervi trigemini, RHmV; nucleus motorius nervi facialis, RHmVII; nucleus motorius nervi glossopharyngei et nervi vagi, RHmIX & X; Fig. 14–17, E, F). In these nuclei, no expression was observed for either *VGluT* or *GAD1* (Fig. 14–17, B, C). Nevertheless, all dopamine-related genes except for *DATx2* were also expressed in this region (Fig. 14–17, G–L). We also found a dense AchE staining signal and *ChAT* expression in a medial part of the isthmic region (Fig. 14–15, E, F), on which we discuss in the next section. Interestingly, *SerT* expression was observed in the RHmV (Fig. 14–15, D).

### Special remarks on the cerebellum-like area and NCvl

Based on the molecular profiling described above, we conducted further examinations of two brain regions that have been involved in the discussion of homology between hagfish and other vertebrates: the cerebellum-like area and NCvl.

#### Cerebellum-like area

It has been suggested that hagfish lacks the proper cerebellum. Still, Larsell (1947) described a cluster of neurons in the isthmic region, and suggested that this brain region is the primordial cerebellum. Furthermore, Kusunoki et al. (1982) reported that strong AchE signals are observed in the neuronal somata and fibers in this region, dorsal to the fibrae arcuatae internae (fai). Based on the fact that AChE signal is also observed in some neuronal somata of the teleost cerebellum (Contestabile, A. & Zannoni, 1975; Contestabile, Antonio, 1975), they suggested that these AChE signals support the existence of the primordial cerebellum in hagfish (Kusunoki et al., 1982).

Our analysis confirmed a cluster of neurons dorsal to fai, and that AChE signals (both neuronal somata and fibers) in this region as well as some neurons in fai (Fig. 18A, E). The expression analysis showed that some of these neurons express ChAT, suggesting that these are indeed cholinergic neurons (Fig. 18F).

**Figure 18.**
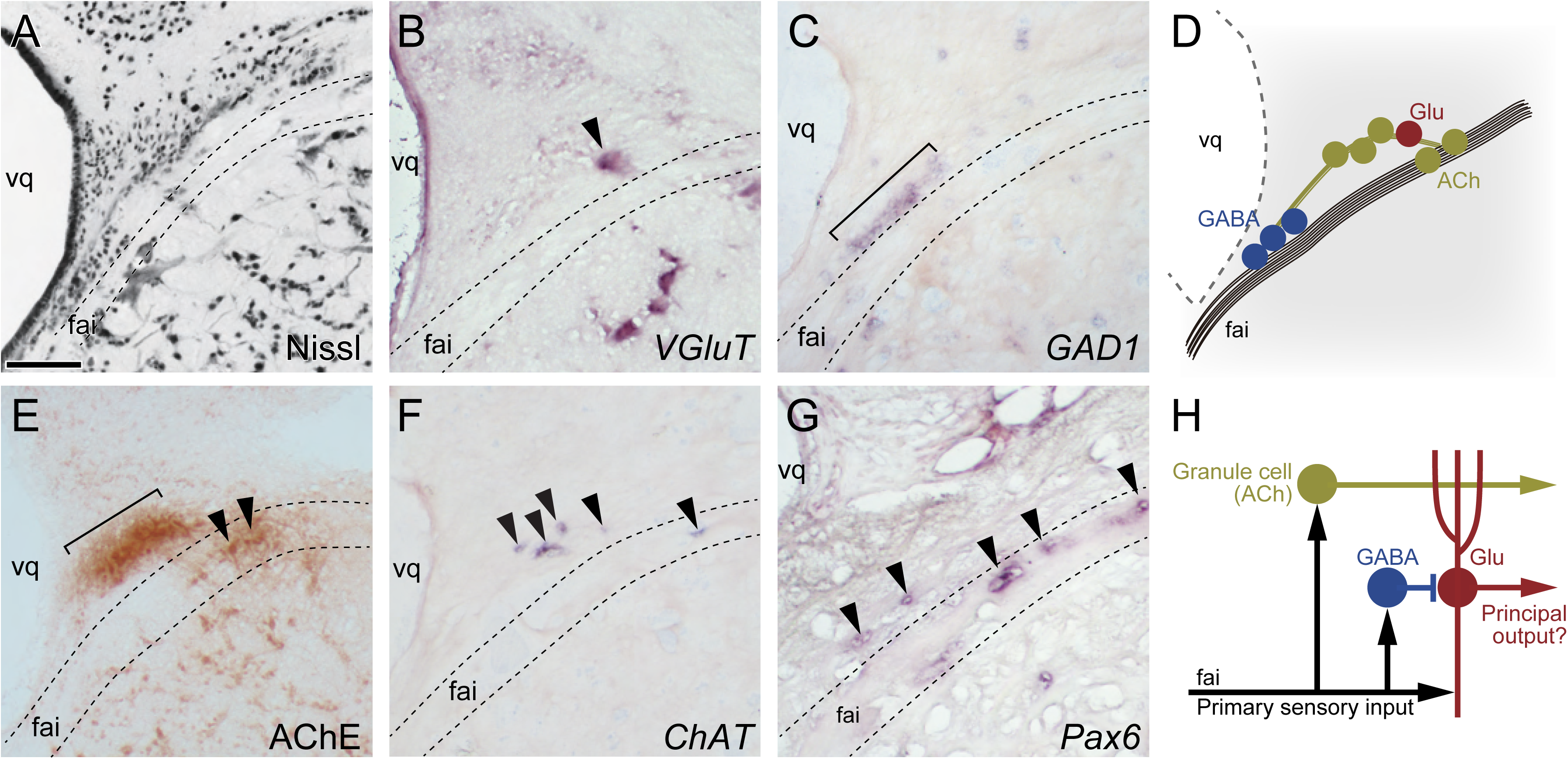
Molecular profiling of the cerebellum-like area. (A–C, E–G) Histological and gene expression analysis in transversal sections: Nissl staining (A), *VGluT* (B), *GAD1* (C), AChE staining (E), *ChAT* (F), *Pax6* (G). (D) Schematic summary of the molecular profiling. (H) Schematic image of the cerebellum-like neural circuit. Scale bar: 100 μm in (A) for (A–C, E–G).

*Pax6* is known to be expressed in mouse embryonic and adult cerebellum (Duan et al., 2013). Previously, Sugahara et al. (2021) isolated hagfish *Pax6* and found that it is expressed in the rhombencephalic lip (i.e., the cerebellum progenitor) of the hagfish embryo. Here, we confirmed *Pax6* expression in some neurons dorsal to fai, as well as those in this neuronal tract (Fig. 18G).

Furthermore, we found that *VGluT* is expressed in a few large neurons dorsal to fai, and GAD1-positive cells are located medially to these *VGluT*-positive cells in a clustered manner (Fig. 18B, C).

#### NCvl

NCvl is a part of the central prosencephalic complex and bilaterally located in the core of the forebrain. This brain region is recognized as an anterocaudally elongated nucleus or dense cluster of neuronal somata (Fig, 19A, B). The three-dimensional reconstruction revealed that the anterior NCvl is sandwiched between the vlt and vld (Fig, 19A). Notably, the vlt extended itself to the ventral side of the anterior NCvl, implying that vlt and vld are primarily formed continuously in early development (Fig, 19A, dashed arrows; Wicht & Nieuwenhuys, 1998), covering the ventral half of the anterior NCvl, and then secondary segregated from each other.

NCvl has been arguably suggested to be homologous to the hippocampus (Holmgren, 1919). To examine this hypothesis, we first focused on the transcription factor gene *Tbr1*, which is known to be expressed in the mouse hippocampus (Bulfone et al., 1995; Medina et al., 2004). We thus isolated hagfish *Tbr1* and conducted expression analysis for phylogenetic analysis (see Suppl. Fig. 3A). As a result, we found that hagfish *Tbr1* is expressed in the NCvl as well as NCm (Fig, 19C).

Furthermore, we also confirmed that *GAD1* is expressed in some NCvl neurons, and that dense cholinergic fibers are distributed in this region (Fig, 19D, E). As the hippocampus contains GABAergic neurons and receives cholinergic inputs (Pelkey et al., 2017), these results further support the homology between NCvl and the hippocampus. To confirm that NCvl receives cholinergic inputs, we identified hagfish muscarinic acetylcholine receptors (mAChRs; for phylogenetic analysis, see Suppl. Fig. 3B) and confirmed that hagfish *mAChR1×1* is expressed in NCvl but not in NCm (Fig, 19F).

## Discussion

In this study, we first clarified the three-dimensional structure of the hagfish brain (Fig. 1). As the brain regions are closely packed in hagfish, previous manual drawings have only provided a limited understanding. Our computer graphics-based approach offers precise topographical information for future neuroanatomical and neurophysiological studies.

Having established the detailed brain architecture, we next examined expression patterns of genetic markers for major neurotransmitters (Fig. 2–17, summarized in Table 2). As previous research has relied on immunohistochemistry, molecular information has been restricted by the availability and cross-reactivity of the antibodies, of which specificity is not guaranteed. By applying *in situ* hybridization technique, gene-specific signals of various molecules were obtained in this study.

In particular, the chemoarchitecture of glutamate and GABA has not been studied in the hagfish brain because of the lack of the appropriate antibodies (Wicht & Northcutt, 1992). Glutamatergic neurons are the most common excitatory neurons in vertebrates (Martinez-Lozada & Ortega, 2023). Also in hagfish, *VGluT*-positive cells were observed in different brain regions, including the pallium, thalamus, and tectal deep layer. In contrast, GABAergic neurons chiefly act as inhibitory neurons. Hagfish *GAD1*-positive cells were also found in several brain regions, particularly notably in the striatum. As the striatum of other vertebrates is known to contain many GABAergic neurons (lamprey, Lamanna et al., 2023; shark, Quintana-Urzainqui et al., 2012; zebrafish, Filippi et al., 2014; mouse, Lein et al., 2007), this result support the homology of the striatum among these animals.

AChE staining was widely used to detect cholinergic neurons but its specificity has been doubted. Co-localization assay of AChE staining and ChAT immunohistochemistry showed that there are many AChE-positive but ChAT-negative cells in rat cerebrum, suggesting that ChAT is a more precise marker for cholinergic neurons than acetylcholinesterase (Levey et al., 1983). Additionally in our analysis, distribution patterns of the strong AChE staining signals and *ChAT*-positive cells were generally consistent, but weak AChE signals were more broadly observed. This result supports the high specificity of the *in situ* hybridization signals.

Kadota (1991) reported that hagfish serotonergic neurons are located in the raphe nucleus and the infundibular region, with serotonergic fibers widely distributed in the forebrain. In this study, we found *SerT*-positive neurons not only in the raphe nucleus and the infundibular region, but also in several forebrain regions. This result suggests these forebrain regions contain not only serotonergic fibers but also somata, and how the immunohistochemical fiber signals masked somatic signals in the previous research.

### Diversity of the dopaminergic system

In our gene expression analysis of major neurotransmitter markers, the most notable finding is the broad distribution of the hagfish dopaminergic system. We found that two distinct dopaminergic markers (*TH* and *DATx1*) are expressed in various brain regions throughout the hagfish brain (Fig. 3–17G, H). It is widely known that vertebrate dopaminergic neurons are predominantly restricted to specific brain regions: the olfactory bulb, the basal diencephalon (including hypothalamus), and midbrain tegmentum (Yamamoto & Vernier, 2011). In lungfish, however, TH-positive neurons are reported to be distributed in other brain regions, including the pallium (medial, dorsal, and lateral), preoptic area, and prethalamus (López & González, 2017). Pallial and subpallial TH-positive neurons are described in chondrichthyans as well (Carrera et al., 2012; Meredith & Smeets, 1987; Northcutt, R. G. et al., 1988; Stuesse et al., 1990, 1991). Taken together, it is suggested that the dopaminergic system once played a broader role in early vertebrate evolution and was secondarily restricted in its distribution.

Additionally, our phylogenetic and synteny analyses highlight another unique aspect of the hagfish dopaminergic system. First, we found two dopamine/norepinephrine transporter candidates: DATx1 and DATx2. Our phylogenetic analysis showed affinity between hagfish DATx2 and gnathostome DAT, and DATx1 formed a sister group to all other vertebrate DAT and NET (Fig. 2D). Nevertheless, microsynteny analysis indicates similarity between hagfish DATx1 and gnathostome DAT, and between hagfish DATx2 and gnathostome NET, respectively (Suppl. Fig. 1). As we observed no significant *DATx2* expression (Fig. 3–17D), DATx1 appears to function as the sole dopamine (or possibly both dopamine and norepinephrine) transporter in the hagfish brain. Notably, Lovell et al. (2015) reported that the *DAT* gene has been lost in the sauropsids (birds and reptiles), and that its function is compensated for by the expression of the *NET* gene. These findings suggest that dynamic and flexible modifications of the molecular mechanisms have been involved in the evolution of the dopaminergic/norepinephrinergic neural circuits in vertebrates.

Next, we found two distinct dopamine D1 receptors (Drd1×1 and Drd1×2) in hagfish, while lampreys have only one receptor. Our phylogenetic analysis showed that the hagfish Drd1×1 and Drd1×2 forms clusters with gnathostome Drd1A and Drd1C, respectively, although the bootstrap values are not high enough (Fig. 2F). As Yamamoto et al. (2015) suggested that the common ancestor of the vertebrates had two types of D1 receptors: D1A/B and D1C/E. If this is the case, our findings suggest that these two types are conserved in hagfish but one of them (presumably D1A/B) has been lost in lampreys. In contrast, our genomic and phylogenetic analyses suggest that hagfish have one D4 and no D2 receptors, while lampreys have two D4 (Drd4 and Drd4-like) and one D2 (Fig. 2G, Suppl. Fig. 2). Our phylogenetic analysis indicated that hagfish Drd4 and lamprey Drd4-like form a sister group with gnathostome Drd4rs, while lamprey D4 shows affinity to gnathostome D4 (Fig. 2G). This result suggests that the common ancestor of the vertebrates had two types of D4, contrary to Yamamoto et al. (2015), and that hagfish had lost D2 and lamprey/gnathostome D4-equivalent genes. Accordingly, both hagfish and lampreys appear to have experienced more complex evolution of the D2/4 family than previously thought (Perez-Fernandez et al., 2016; e.g., Yamamoto et al., 2015).

### Evolutionary origin of the cerebellum

It is generally accepted that cyclostomes lack the proper cerebellum (Kebschull et al., 2020; Lamanna et al., 2023; Lannoo & Hawkes, 1997; Wicht & Nieuwenhuys, 1998). However, Sugahara et al. (2021) demonstrated embryonic evidence of molecular patterning for the differentiation of the cerebellar structures in hagfish and lampreys. Moreover, Kusunoki et al. (1982) suggested that hagfish have AChE-positive granule cells. To thus elucidate the evolutionary origin of the cerebellum, cerebellum-related features of the hagfish brain are required to be examined.

Bell et al. (2008) argued that most vertebrates have both the cerebellum and structures that are architecturally similar to the cerebellum, and that cerebellum-like structures may have evolved before the cerebellum itself. For example, medial and dorsal octavolateral nuclei (MON and DON, respectively) exhibits cerebellum-like neuroarchitecture with an exception that principal output is not inhibitory as Purkinje cells but excitatory (Bell et al., 2008; Montgomery et al., 2012; Striedter & Northcutt, 2019).

This study provides a first molecular profiling of the cerebellum-like area in hagfish (summarized in Fig. 19H). We confirmed a cluster of AChE- and *ChAT*-positive cells dorsal to fai, which also contains some of these cells as well (Fig. 19E, F). Just ventromedial to this cluster, on the dorsal surface of fai, a series of *GAD1*-positive cells were observed (Fig. 19C). In gnathostomes, *Pax6* is known to be expressed in granule cell layer (Yamasaki et al., 2001). We found some *Pax6*-positive cells in and dorsal to fai (Fig. 19G). These results suggest that cholinergic and GABAergic neurons are involved in the hagfish cerebellum-like neuroarchitecture, possibly equivalent to excitatory granule cells and inhibitory Golgi cells in gnathostome cerebellum, respectively. In addition, we found some VGluT-positive cells nearby the cluster of AChE- and *ChAT*-positive cells; these cells appear to correspond to the principal output neurons.

**Figure 19.**
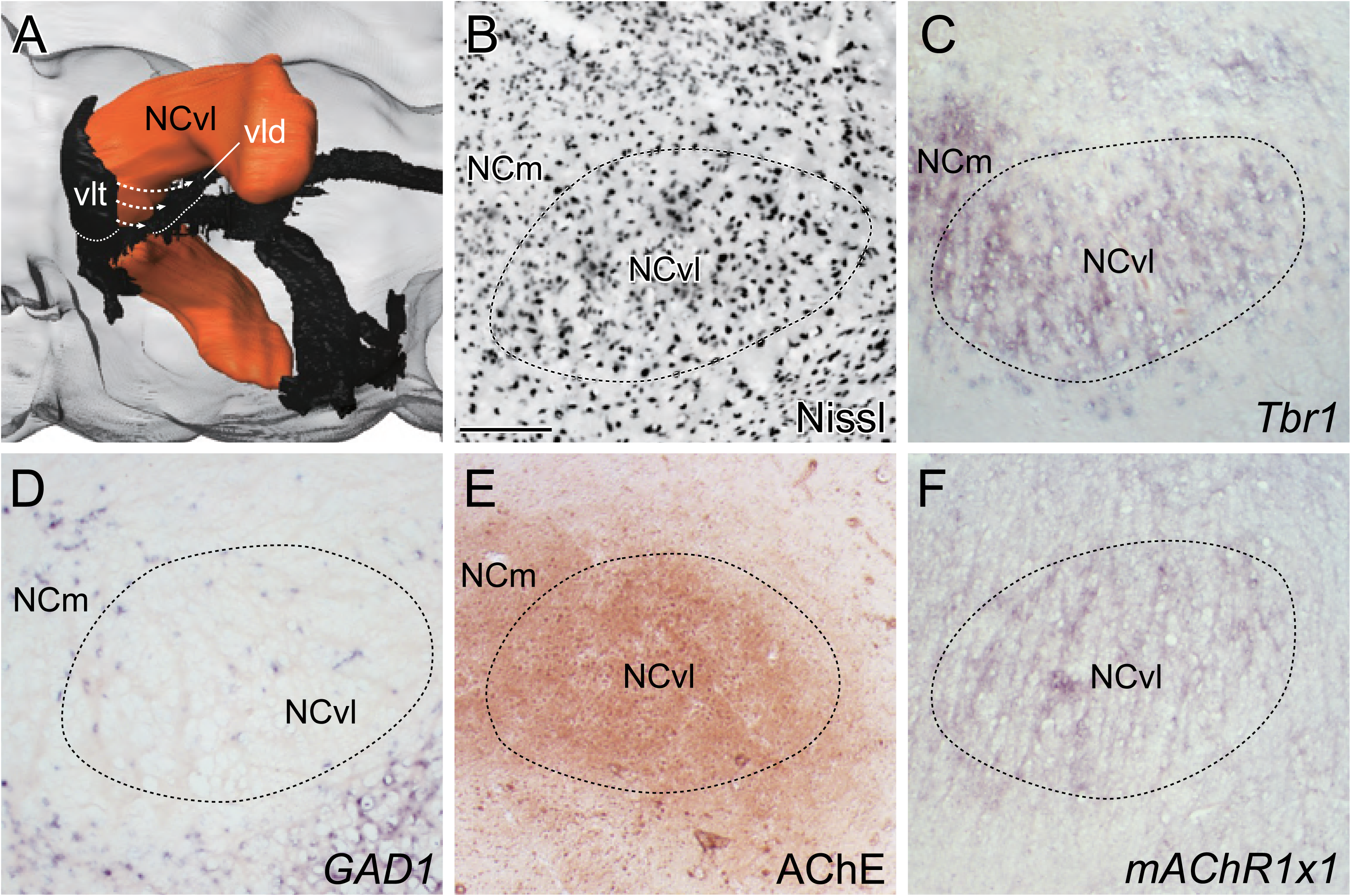
Molecular profiling of the NCvl. (A) Three-dimensional reconstruction of the NCvl with vlt and vld in ventrolateral view. (B–F) Histological and gene expression analysis in transversal sections: Nissl staining (B), *Tbr1* (C), *GAD1* (D), AChE staining (E), *mAchR1×1* (G). Scale bar: 200 μm in (B) for (B–F).

Our results suggest that the cerebellum-like area in hagfish shows a neural circuit similar to that of the cerebellum-like MON/DON in jawed vertebrates, supporting the idea that the common ancestor of the vertebrates had the cerebellum-like neuroarchitecture prior to the acquisition of the proper cerebellum in the gnathostome lineage. Further research including co-expression and neurocircuitry analyses will help to elucidate the evolutionary origin of the cerebellum at the neuronal level.

### Evolution and development of the forebrain structures

The central part of the hagfish forebrain is occupied by an enigmatic structure: the central prosencephalic complex. Two major hypotheses have been proposed regarding which gnathostome brain region corresponds to this structure: the hippocampus or prethalamic eminence (PThE, also known as the thalamic eminence).

Based on the cellular morphology, Holmgren (1919) compared the central prosencephalic complex to the “primordium hippocampi” (Johnston, 1912) or the medial pallium of the lamprey. Nonetheless, these structures are placed at different positions in the forebrain; the former occupies the central part covered by the pallium and striatum (Fig. 1E, F), while the latter is situated in the dorsomedial part (see Fig. 20). Holmgren (1919) tried to explain this topological disparity with his hyperinversion theory, where the lateral ventricle corresponds to Wicht & Northcutt’s (1992) P3 region and the pallium develops from ventral to dorsal, covering the central prosencephalic complex (Wicht & Northcutt, 1992).

**Figure 20.**
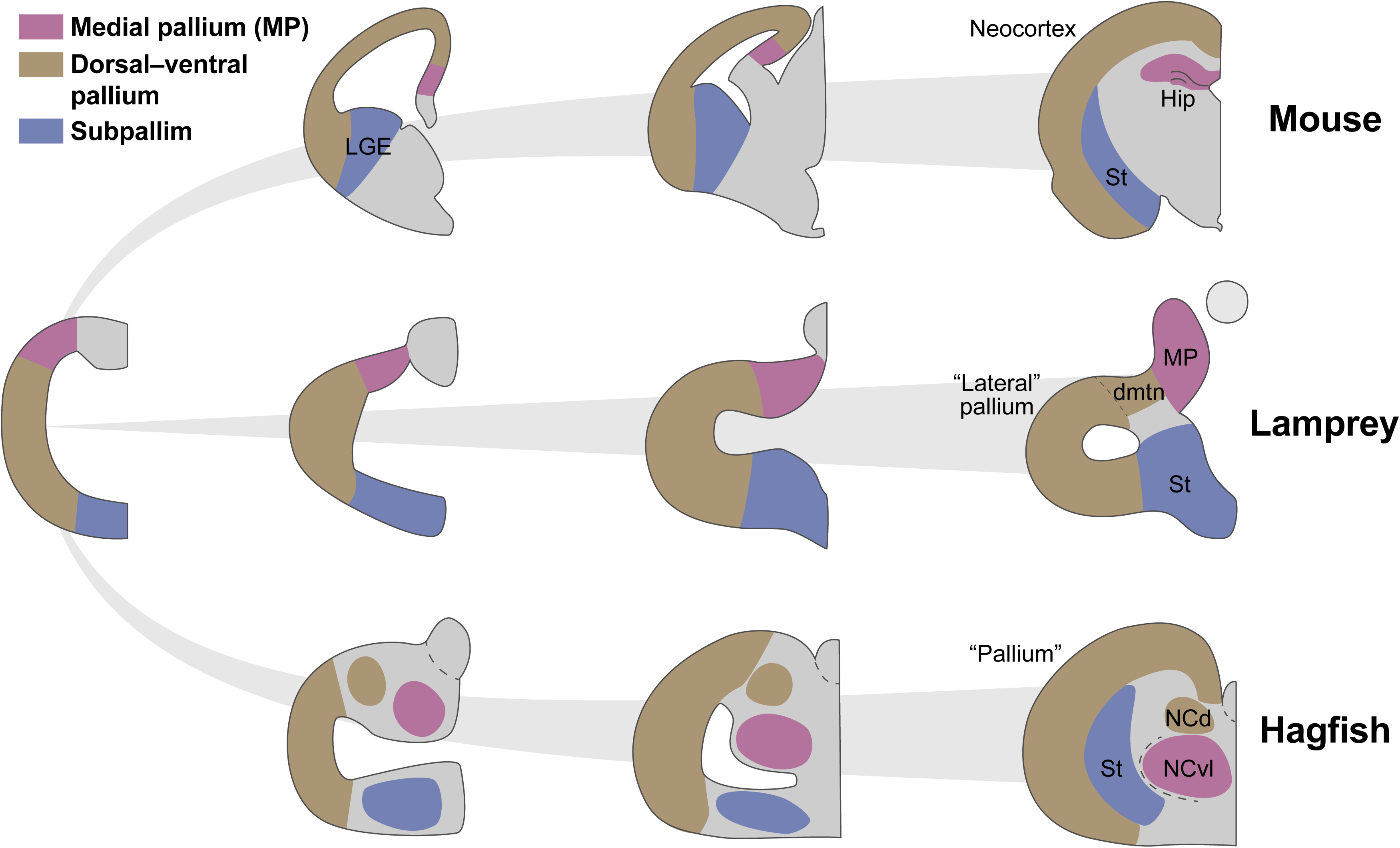
Schematic image of the forebrain structure development. The cross sections of the mouse (top), lamprey (middle), and hagfish (bottom) forebrains are shown from early (left) to later (right) stages.

Against this idea, Wicht & Northcutt (1992) argued that the central prosencephalic complex is not homologous to hippocampus but to PThE, based on several grounds. They first pointed out that it is located caudal to the “vestiges of the lateral ventricle” and can be traced caudally well into diencephalic areas. However, our analysis showed that it is located between vlt and vld, of which only the former one was recognized by the authors. Therefore, the topological organization still retains the possibility that this structure is derived from the medial pallium as discussed below.

Hodology, or neural connectivity, was another criterion employed by Wicht & Northcutt (1992, 1998). They noted that none of the nuclei of the central prosencephalic complex receives any ascending thalamic fibers in hagfish, while the medial pallium is the main pallial target of ascending thalamic projections in many gnathostomes (Northcutt, 1995) and in lampreys (Northcutt & Wicht, 1997; Polenova & Vesselkin, 1993). Amemiya & Northcutt (1996) contradicts this statement, showing that tracer injection to the NCm and NCvl results in retrogradely labeled neuronal somata in THsh. However, more detailed analyses are required as they did not inject tracer to NCm and NCvl separately, coupled with uncertainties in their interpretation of brain regions due to lack of three-dimensional brain atlas. Additionally, Wicht & Northcutt (1998) argued that some neural connections of NCm and NCvl (connections with the preoptic area and habenula; Amemiya & Northcutt, 1996) match those of the PThE of axolotols described by Krug et al. (1993). Still, the gnathostome hippocampus has shown to have connections with the corresponding regions (Cenquizca & Swanson, 2006; Goutagny et al., 2013). The connections with the preoptic area and habenula thus cannot be the determinative factor for the homology between NCvl and PThE. In addition, AChE-positive fibers and *mAChR1×1*-expressing cells in NCvl shown in this study (Fig. 19D, E) strongly suggest that this region receives cholinergic inputs similar to the lamprey medial pallium (Pombal et al., 2001) and the gnathostome hippocampus (Teles-Grilo Ruivo & Mellor, 2013).

The expression pattern of *Tbr1*, shown in this study, provides a hint to resolve the current disagreement, since this transcription factor is known to be expressed not only in the hippocampus but also in the PThE (Alonso et al., 2020). As *Tbr1* was expressed both in NCm and NCvl but *mAChR1×1*-positive cells were observed in the latter (Fig. 19C, E), the former and the latter may correspond to PThE and the hippocampus, respectively. Moreover, the other subnucleus of the central prosencephalic complex, NCd, is reported to receive dense secondary olfactory projections from the olfactory bulb (Wicht & Northcutt, 1993). Recently, Suryanarayana et al. (2021) demonstrated that a lamprey brain region between the lateral and dorsal pallia, called the dorsomedial telencephalic nucleus (dmtn), receives secondary olfactory inputs from tuft-like neurons in the olfactory bulb. These studies suggest the affinity between the hagfish NCd and the lamprey dmtn.

Taken together, we propose that NCvl, NCm, and NCd corresponds the hippocampus, the lamprey dmtn, and the PThE, respectively, and that these subnuclei of central prosencephalic complex migrate from dorsal part to the central core of the forebrain due to the large expansion of the dorsolateral pallium (Fig. 20). This model may be regarded as a revised version of Holmgren’s (1919) hyperinversion theory, where the position of the lateral ventricle is corrected to be between NCvl and striatum, according to our three-dimensional reconstruction. Although single cell transcriptome analysis of the lamprey brain suggests that the excitatory neurons in its medial pallium express genetic markers not for pallial but for pre-thalamic excitatory neurons (Lamanna et al., 2023), further developmental and special transcriptomic studies are required to determine the homology of forebrain structures between hagfish, lampreys, and jawed vertebrates.

### Limitations and future perspectives

In this study, we have conducted three-dimensional reconstruction and gene expression analysis to establish a modern brain atlas of hagfish. Still, there are also several limitations, some of which have already been mentioned above. First, only a limited number of genes were examined here. Further gene expression analyses including spatial transcriptomics will help to provide a more comprehensive understanding of the gene expression patterns in specific brain regions and neurons. Second, some of our gene expression results are not consistent with previous reports using immunohistochemistry. This should be examined if it is due to the protein synthesis and localization processes or due to antibody epitope reactivity. Third, the development of the hagfish brain regions remains to be clarified in order to allow comparison with that of lampreys and jawed vertebrates. Last, further research on neural connections between brain regions would be facilitated by combining fluorescent tracer injections with the three-dimensional brain model presented in this study. Despite these limitations, our results provide a promising foundation for contemporary hagfish neurobiology, establishing this animal as a resurgent model for investigating the evolutionary origin of the vertebrate brain.

## Acknowledgment

We thank Susumu Hatanaka (Shinsho Maru; Fujisawa, Kanagawa, Japan) for providing the *E. burgeri* specimens. This work was supported by the Grant-in-Aid for the Japan Society for the Promotion of Science (JSPS; Grant Numbers JP20K15855, JP22K15164, JP24K09556, and JP24H01538 to D.G.S.), NIBB Collaborative Research Project for Integrative Imaging (21-420, 22NIBB520, 23NIBB511, 24NIBB519, and 25NIBB516 to D.G.S.), and by the Sasakawa Scientific Research Grant from The Japan Science Society (Grant Number 2024-5028 to R.H.).

## Supplemental materials

**Supplemental Figure 1.** Microsynteny analysis of *DAT*/*NET*. For the details of the flanking genes, see Supplementary Table 2.

**Supplemental Figure 2.** Microsynteny analysis of *Drd2* and its flanking genes. For the details of these genes, see Supplementary Table 3.

**Supplemental Figure 3.** Phylogenetic trees of Tbr1 and mAChRs, based on amino acid sequences. (A) Tbr1, (B) mAChRs. The numbers described on some nodes indicate exact bootstrap values. Scale bar represents the number of amino acid substitutions per site.

**Supplemental** Table 1. Nomenclature of the hagfish brain regions.

**Supplemental** Table 2. Details of the flanking genes to DAT/NET for microsynteny analysis.

**Supplemental** Table 3. Details *Drd2* and its flanking genes for microsynteny analysis.

## Abbreviations

A: anterior
BO: bulbus olfactorius
BOci: bulbus olfactorius, stratum cellulare internum
BOg: bulbus olfactorius, stratum glomerulosum
BOm: bulbus olfactorius, stratum mitrale
BOn: bulbus olfactorius, stratum nervosum
BOp: bulbus olfactorius, stratum periglomerulosum
D: dorsal
DIE: diencephalon
HA: habenula
HAcde: habenula, corpus habenulae dextrae
HAcsi: habenula, corpus habenulae sinistrae
HAl: habenula, nucleus lateralis
HAv: habenula, nucleus ventralis
HY: hypothalamus
HYinf: hypothalamus, nucleus infundibularis
HYnhyp: hypothalamus, neurohypophysis
MES: mesenceobalon
NC: nucleus centralis
NCd: nucleus centralis procencephali, pars dorsalis
NCm: nucleus centralis procencephali, pars medialis
NCvl: nucleus centralis procencephali, pars ventrolateralis
P: posterior
P1–5: pallium layer 1–5
PO: preoptic area
POd: area praeoptica, nucleus dorsalis
POe: area praeoptica, nucleus externus
POim: area praeoptica, nucleus intermedius
POp: area praeoptica, nucleus internus or periventricularis
RH: rhombencephalon
RH“A”: rhombencephalon, nucleus “A” of Kusunoki et al. (1982)
RH“B”: rhombencephalon, nucleus “B” of Kusunoki et al. (1982)
RHdc: rhombencephalon, nuclei columnae dorsalis
RHmV: rhombencephalon, nucleus motorius nervi trigemini
RHmVII: rhombencephalon, nucleus motorius nervi facialis
RHmIX&X: rhombencephalon, nucleus motorius nervi glossopharyngei et nervi vagi
RHoc: rhombencephalon, area octavolateralis
RHra: rhombencephalon, nuclei raphe
RHrdV: rhombencephalon, nucleus radicis descendens nervi trigemini
RHri: rhombencephalon, formatio reticularis, pars inferior
RH“rl”: rhombencephalon, rostrolateral part of the reticular formation of Kusunoki et al. (1982)
RH“rm”: rhombencephalon, frostromedial part of the reticular formation of Kusunoki et al. (1982)
RHrm: rhombencephalon, formatio reticularis, pars medialis
RHrs: rhombencephalon, formatio reticularis, pars superior
RHv: rhombencephalon, area ventralis
Se: septum
St: striatum
SY: synencephalon
SYcp: synencephalon, nucleus commissurae posteriois
SYflm: synencephalon, nucleus fasciculi longitudinalis medialis
SYinp: synencephalon, nucleus interpeduncularis
SYpt: synencephalon, nucleus praetectalis
TEL: telencephalon
TEG: tegmentum
TEGv: nucleus ventralis tegmentalis
TH: thalamus
THa: thalamus, nucelus anterior
THdi: thalamus, nucelus diffusus
THe: thalamus, nucleus externus
THi: thalamus, nucleus internus
THico: thalamus, nucleus intracommissuralis
THpco: thalamus, nucleus paracommissuralis
THsh: thalamus, nucelus subhabenularis
THt: thalamus, nucleus triangularis
TM: tectum mesencephali
TMscf: tectum mesencephali, stratum cellulare et fibrosum
TMsm: tectum mesencephali, stratum marginale
TMsp: tectum mesencephali, stratum periventriculare
TP: tuberculum posterior
TPl: tuberculum posterius, nucleus lateralis
TPm: tuberculum posterius, nucleus medialis
V: ventral

