## Supplementary figures and images for "Three-dimensional and molecular brain atlas of the hagfish reveals the evolutionary origin and early diversification of the vertebrate brain"

### Supplemental Fig. 1

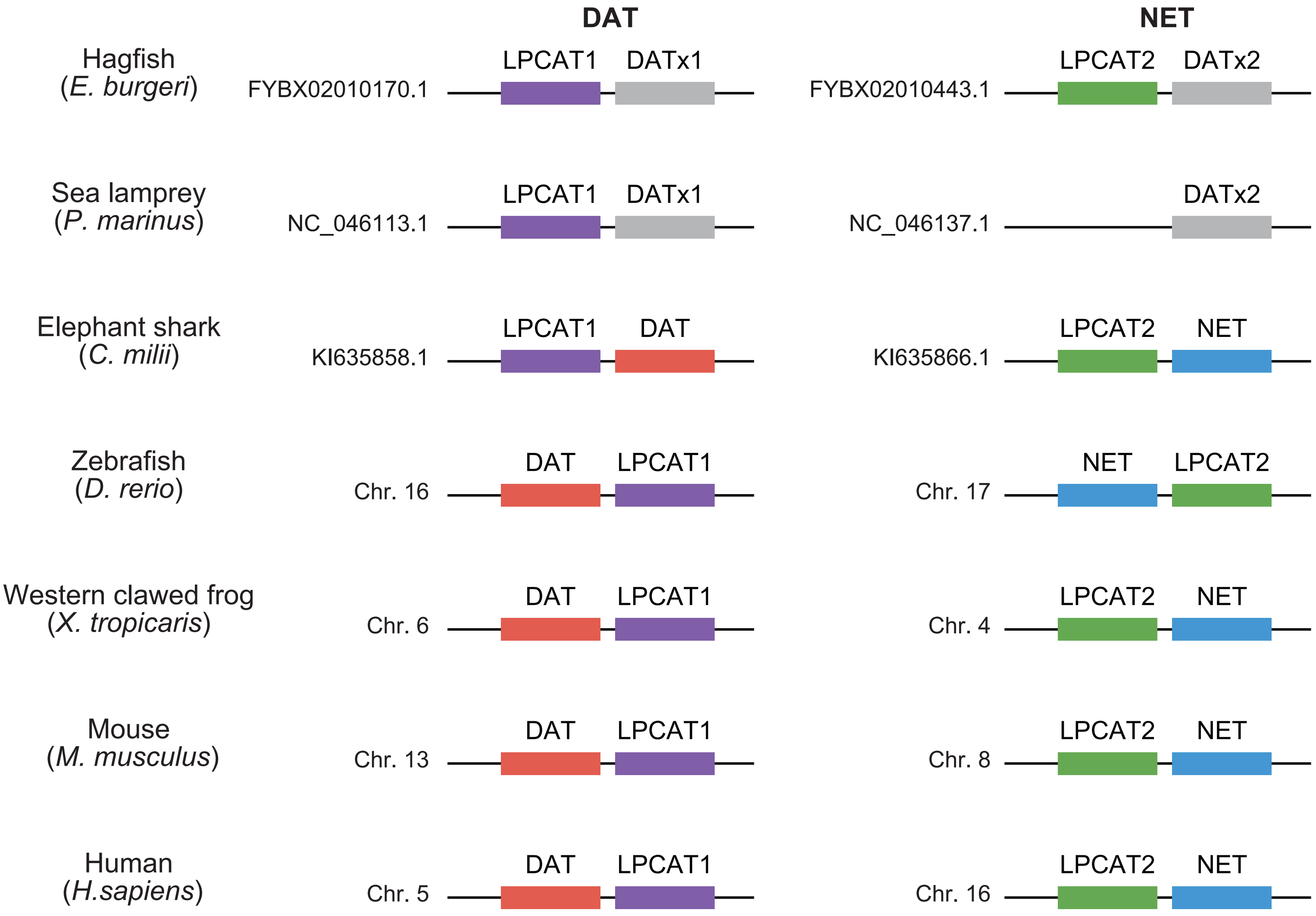

### Supplemental Fig. 2

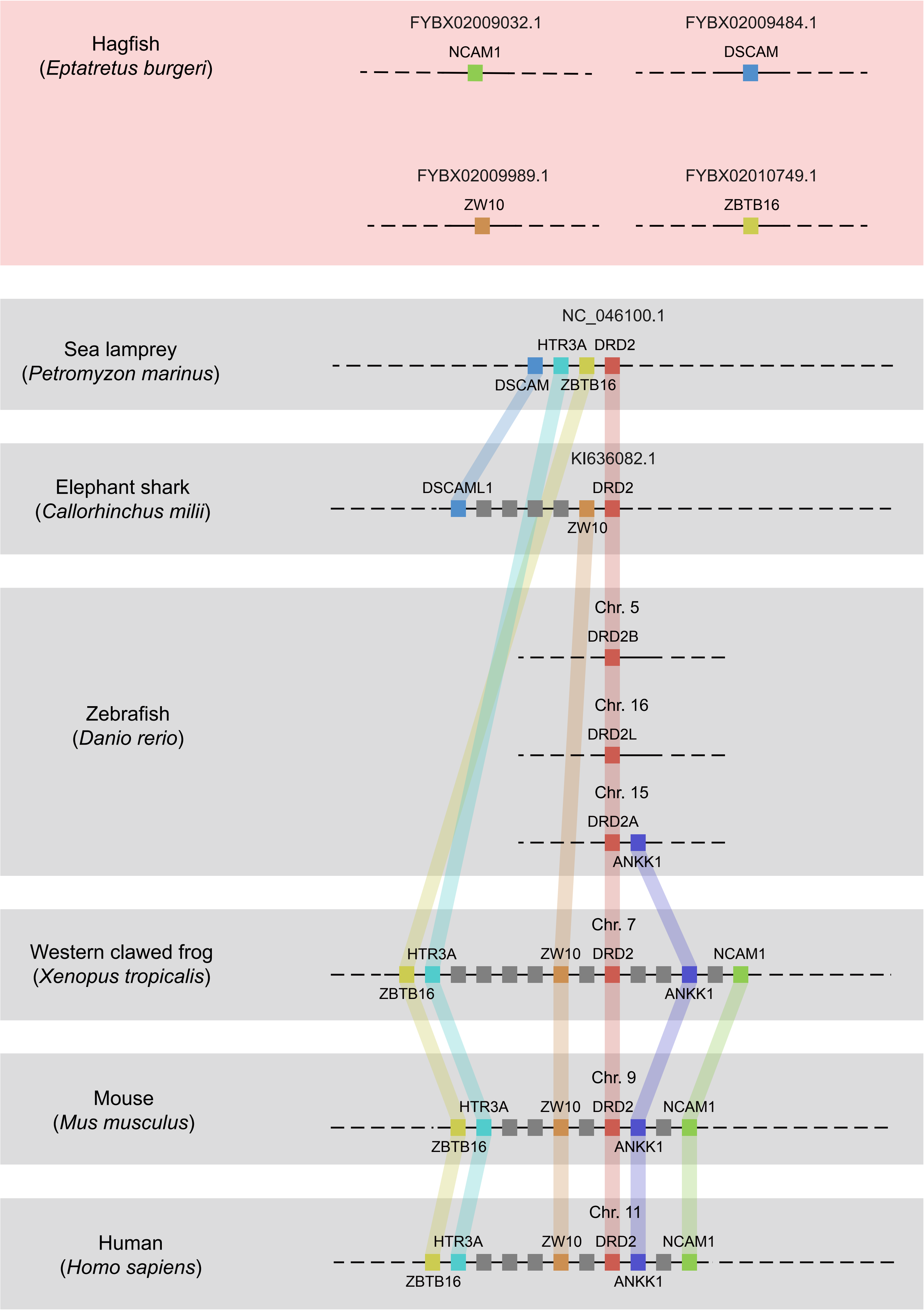

### Supplemental Fig. 3

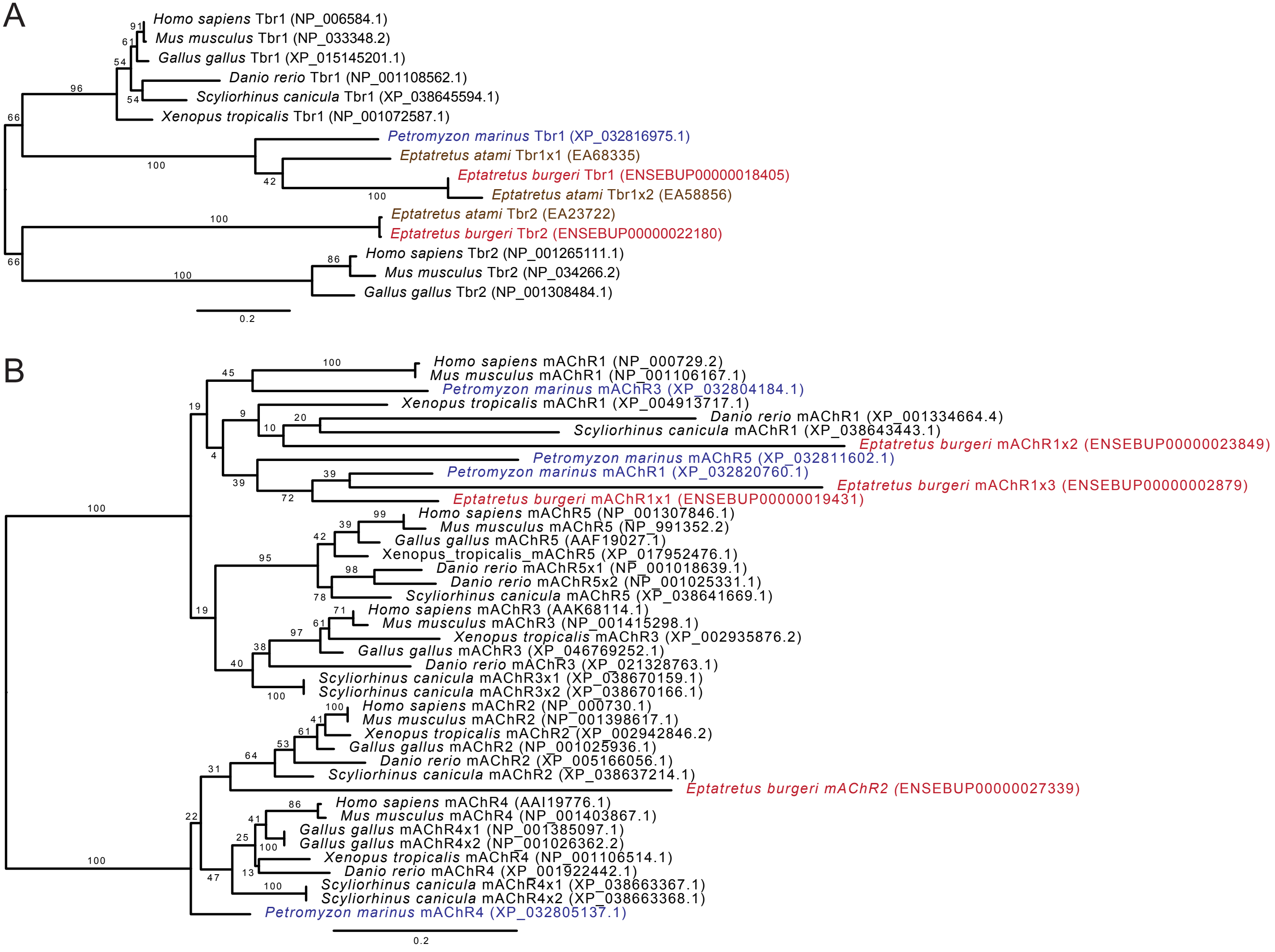
